# The Outer Kinetochore Proteins KNL-1 and Ndc80 complex are Required to Pattern the Central Nervous System

**DOI:** 10.1101/2024.03.27.586904

**Authors:** Vasileios R. Ouzounidis, Mattie Green, Charlotte de Ceuninck van Capelle, Clara Gebhardt, Helena Crellin, Cameron Finlayson, Bram Prevo, Dhanya K. Cheerambathur

**Affiliations:** Wellcome Centre for Cell Biology & Institute of Cell Biology, School of Biological Sciences, The University of Edinburgh, Edinburgh, EH9 3BF, UK

## Abstract

The KMN (Knl1/Mis12/Ndc80) network at the kinetochore, primarily known for its role in chromosome segregation, has been shown to be repurposed during neurodevelopment. Here, we investigate the underlying neuronal mechanism and show that the KMN network is essential to establish the proper axonal organization within the *C. elegans* head nervous system. Post-mitotic degradation of KNL-1, which acts as a scaffold for signaling and has microtubule-binding activities at the kinetochore, led to disorganized ganglia and aberrant placement and organization of axons in the nerve ring - an interconnected axonal network. Through gene-replacement approaches, we demonstrate that the signaling motifs within KNL-1, responsible for recruiting the protein phosphatase 1, and activating the spindle assembly checkpoint are required for neurodevelopment. Interestingly, while the microtubule-binding activity is crucial to KMN’s neuronal function, microtubule dynamics and organization were unaffected in the absence of KNL-1. Instead, the NDC-80 microtubule-binding mutant displayed notable defects in axon bundling during nerve ring formation, indicating its role in facilitating axon-axon contacts. Overall, these findings provide evidence for a non-canonical role for the KMN network in shaping the structure and connectivity of the nervous system in *C. elegans* during brain development.

## INTRODUCTION

The central nervous system (CNS) relies on the precise wiring of neurons to regulate and coordinate an organism’s behaviour and responses to the environment (Kolodkin and Tessier-Lavigne, 2011). The assembly of the CNS initiates during the embryo stage and is a highly orchestrated developmental process that involves a complex interplay of neuronal ganglia, axons, dendrites, and synapses to create a highly intricate three-dimensional structure (Bénard and Hobert, 2009). Despite significant progress in understanding the spatial and developmental cues that direct CNS formation, how neuronal downstream effectors such as cell adhesion molecules and cytoskeletal structures facilitate the assembly and maintenance of the CNS architecture remains unclear.

The tiny roundworm *Caenorhabditis elegans* with its simple and highly stereotypical nervous system, consisting of 302 neurons in the hermaphrodite, is well-suited to study neuronal architecture (White *et al*., 1986). Approximately half of these neurons reside in the worms’ head, where the neuronal cell bodies cluster into ganglia, and their axons converge into a synapse-rich structure termed the nerve ring. The *C. elegans* nerve ring along with the ganglia is considered to be the worm’s brain. The formation of this rudimentary brain occurs during embryonic development through a highly invariant series of orchestrated events (Rapti *et al*., 2017; Moyle *et al*., 2021). During embryogenesis a subset of neurons, the pioneer neurons, extend their axons to build a ring-shaped scaffold that directs the assembly of axons from sensory and interneurons resulting in a dense interconnected nerve ring. As the animal develops, the overall spatial organization of the neuronal cell bodies and the nerve ring remains remarkably stable and persists into adulthood (Bénard and Hobert, 2009). The preservation of this neuronal pattern across individuals and the lifetime of organism suggests the presence of precise mechanisms that govern nervous system architecture.

The neuronal architecture in *C. elegans* is influenced by several molecular factors, including cell adhesion molecules, extracellular matrix proteins and cytoskeletal proteins (Bénard and Hobert, 2009; Barnes *et al*., 2020). For instance, SAX−7, a homologue of the vertebrate L1 family of cell adhesion protein (L1CAM) is involved in axon guidance, dendrite arborization and head ganglia organization (Zallen *et al*., 1999; Sasakura *et al*., 2005; Pocock *et al*., 2008; Bénard *et al*., 2012; Dong *et al*., 2013; Salzberg *et al*., 2013). Additionally, mutations in cytoskeletal proteins such as *unc-44*, the vertebrate ankyrin homologue, and *unc-33,* that encodes for microtubule-binding CRMP protein, also impair axon guidance and fasciculation in several neuronal classes within *C. elegans* (Hedgecock *et al*., 1985; Li *et al*., 1992; Otsuka *et al*., 1995; Zallen *et al*., 1999). Despite these findings, our understanding of the precise molecular identities and mechanisms that facilitate the patterning of the *C. elegans* nervous system remains limited.

Recent studies have revealed essential functions for the evolutionarily conserved chromosome-microtubule coupling machinery, the KMN (Knl1/Mis12/Ndc80-complex) network, during neurodevelopment. Specifically, the KMN proteins are necessary for dendrite extension in *C. elegans*, synapse formation and dendrite regeneration in *Drosophila* and dendrite morphology of rat hippocampal neurons (Cheerambathur *et al*., 2019; Zhao *et al*., 2019; Hertzler *et al*., 2020). The KMN network is best known for its conserved role in mitosis, where KMN complexes facilitate chromosome segregation by forming stable attachments to spindle microtubules (Cheerambathur and Desai, 2014; Cheeseman, 2014; Musacchio and Desai, 2017). During mitosis, KMN ensures the fidelity of chromosome segregation by coordinating mechanical coupling with cell cycle progression and by acting as a scaffold for spindle checkpoint signalling machinery. Within the KMN network, the Ndc80 complex serves as the primary microtubule coupling module (Cheeseman *et al*., 2006; Deluca *et al*., 2006), the Knl1 complex acts as a scaffold for both the Ndc80 complex and signalling machinery, and the Mis12 complex links the KMN network to centromeric chromatin (Musacchio and Desai, 2017). How these discrete functions are repurposed during nervous system development and maintenance across different organisms remains largely unknown.

Here we characterize the role of KMN network in organizing and maintaining the nerve ring axon bundles and head ganglia structure in *C. elegans.* In our prior work on sensory neuronal morphogenesis in *C. elegans*, we found that degradation of KMN components disrupted not only the dendrite formation, but also axon morphology and head ganglion organization (Cheerambathur *et al*., 2019). However, the axonal defects were not characterized and the effect of KMN components on nerve ring structure had not been investigated. In this study, we show that degradation of KNL-1 and Ndc80 complex results in axon placement defects similar to those seen in the absence of the cell adhesion molecule, SAX-7. Furthermore, we show that the axon bundles are defasciculated in the absence of KNL-1. Interestingly, proper axon and ganglia organization within the nerve ring requires both the signalling and microtubule binding functions of KNL-1. Although the global microtubule organization and dynamics of the axon remain unchanged in the absence of KNL-1, the microtubule binding domain of the Ndc80 complex is essential to maintain the integrity of the axonal bundle during its assembly. Thus, our observation suggests that the KMN network play an essential role in the proper development and organization of axons in the central nervous system of *C elegans*.

## RESULTS

### The KMN component KNL-1 is required for axon guidance and positioning during neuronal morphogenesis

To characterize the role of the KMN network during axon formation, we examined the function of KNL-1 in the formation of axons within the amphid neurons, a subset of sensory neurons in the *C. elegans* nerve ring. The amphid, which is the primary olfactosensory organ, consists of twelve bilaterally symmetric neuron pairs, with one dendrite and one axon per neuron (White *et al*., 1986). The amphid neuron has ciliated dendrites that extend anteriorly to the nose of the animal sensing various stimuli while their axons are integrated into the nerve ring, where they form synapses with other neurons (**Figure 1A**). We had previously shown that depletion of KNL-1 from developing amphid neurons, using a tissue-specific protein degradation system, led to both dendrite and axonal morphology defects (Wang *et al*., 2017; Cheerambathur *et al*., 2019). In this system, a GFP nanobody fused to ZIF-1, a SOCS box adapter protein that recruits target proteins to the E3 ubiquitin ligase complex for proteosome mediated degradation, was expressed under the control of the *dyf-7* promoter (**Figure 1B**). This promoter is active throughout the development of most differentiated sensory neurons, including the amphid during embryogenesis (Heiman and Shaham, 2009).

**Figure 1.**
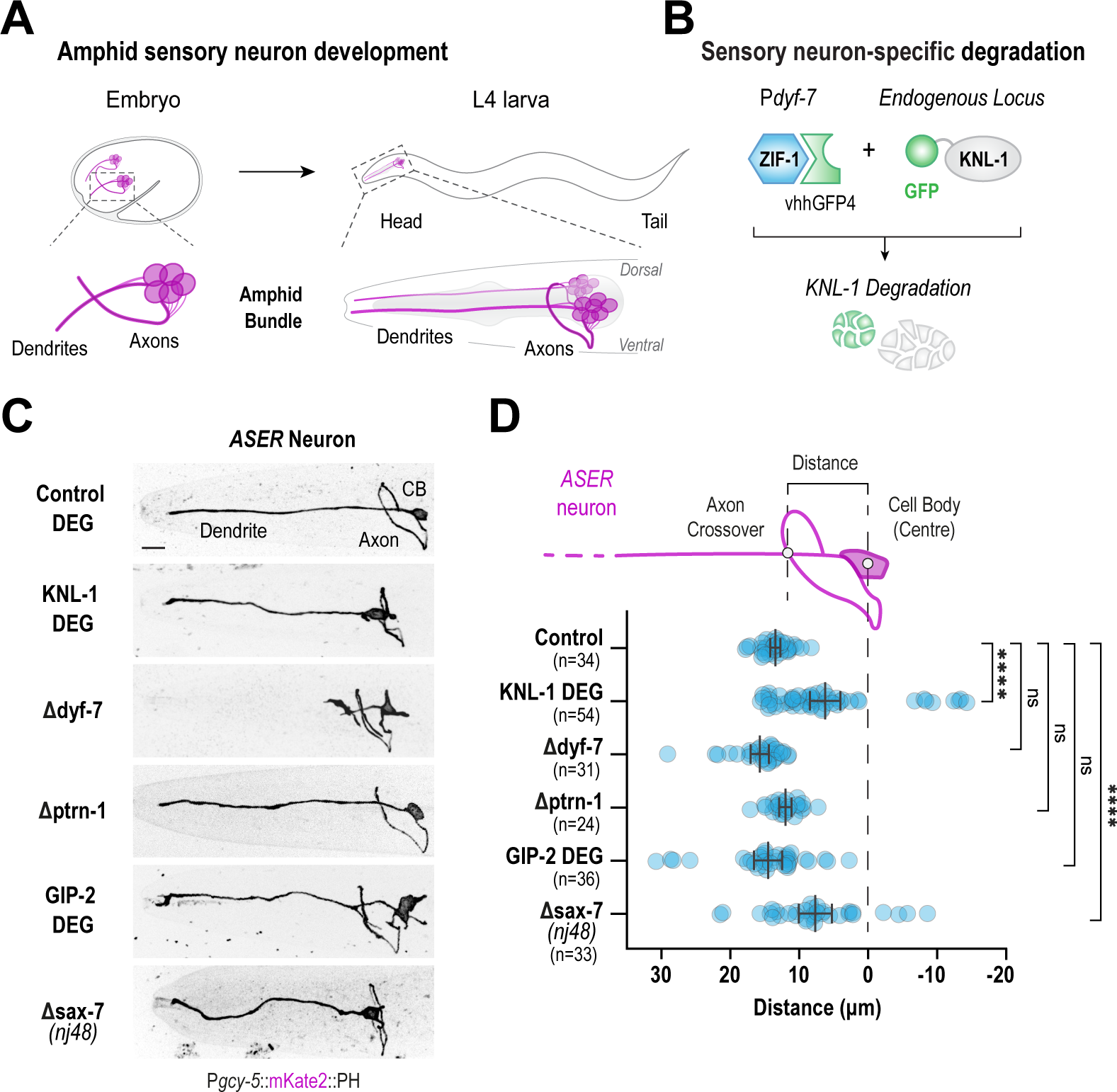
KNL-1 degradation in sensory neurons results in axon placement defects. (A) Schematic of amphid sensory neuron development in *C. elegans*. The cell bodies of the amphid neurons are clustered into ganglia near the pharynx, with each amphid neuron projecting an axon that becomes part of the nerve ring and a single dendrite that extends to the nose. The development of axons within the amphid sensory organ occurs during embryogenesis (left). During larval development, amphid axons increase in length whilst retaining the spatial organization established during embryogenesis. The right cartoon illustrates a lateral view of the mature amphid neurons at the L4 larval stage. (B) Approach used to degrade the in situ GFP::KNL-1 fusion in *dyf-7* positive neurons. The GFP nanobody fused to a SOCS box protein, ZIF-1 which is an E3 ligase adapter, recognizes GFP-tagged proteins and marks them for degradation by the proteosome. (C) Images of ASER amphid neuron morphology at the L4 larval stage. The ASER neuron was labelled using a membrane marker, mKate2::PH (rat PLC1δ pleckstrin homology domain) under the *gcy-5* promoter. The panels represent ASER morphology in the indicated conditions. CB is cell body. Scale bar,10 μm. (D) Schematic of the ASER neuron (top). Plot of distance between the centre of the ASER cell body and the axon-dendrite intersect, as shown in the cartoon above, under different conditions (bottom). n represents the number of animals. Error bars denote 95% confidence interval. **** and ns indicate p<0.0001 and non-significant, respectively.

To further characterize the axonal defects observed upon KNL-1 degradation, we analysed the axonal trajectory of a single amphid neuron, ASER, by using a mKate2::PH (rat PLC-16) marker that labels the membrane and is expressed under an ASER specific promoter, *gcy-5* promoter. In both control and KNL-1 degrader animals, the dendrites of the ASER neuron were extended anteriorly towards the tip of the nose whereas the axonal trajectory of the ASER neuron in the KNL-1 degraded animals significantly differed from that in control animals (**Figure 1C**). In the control, the ASER axon projected anteriorly, relatively to the cell body and displayed the stereotypical trajectory of extending its axonal projection ventrally first and then dorsally within the nerve ring. Although the axons of the ASER neuron in the KNL-1 degrader animals were guided ventrally and entered the nerve ring correctly, the positioning of the axon relatively to the cell body was significantly impacted. Unlike the control animals, where the axons were consistently anterior to the ASER cell body, the axons in KNL-1 degrader animals were misplaced, often found more proximal or posterior to the cell body (**Figure 1C and D**). In control animals the nerve ring entry of the axon was located ∼13.97+/− 1.1μm away from the centre of the ASER cell. However, after KNL-1 degradation, the position of the ASER axon shifted to ∼6.77 +/− 0.3 μm indicating that, in the absence of KNL-1, the axon of the ASER neuron is positioned more posteriorly and closer to the cell body. In our previous study, we had shown that the cell bodies of the sensory neurons were misplaced after KNL-1 degradation. To confirm that axon placement defects were not a result of the displacement of the ASER cell body, we also measured the distance of the axon and cell body position relative to the centre of the terminal pharyngeal bulb situated posterior to the nerve ring. Compared to the control, the ASER axon in the KNL-1 degrader was misplaced posteriorly relatively to the pharynx as well (**Figure S1A**).

We had previously shown that degradation of KNL-1 in sensory neurons impacted their sensory function (Cheerambathur *et al*., 2019). Given that disruptions in sensory activity can affect axon development of amphid neurons, we wanted to ensure that the observed axon positioning defects in KNL-1 degron animals were not a consequence of impaired ASER dendrite structure and function (Peckol *et al*., 1999). To address this, we visualized the ASER neuron organization in the deletion of *dyf-7*. DYF-7 is essential for amphid dendritic extension and animals lacking *dyf-7* lack dendrites that extend to the nose and are thus defective in dendrite structure and signalling (Heiman and Shaham, 2009). Unlike the axonal defects observed in KNL-1 degradation, Δ*dyf-7* animals displayed properly developed axons and the axons were placed anteriorly to the ASER cell body, and the pharynx, at similar distances to that of the control (**Figure 1C, D and S1A**). These results indicate that the placement of axons is not influenced by dendrite morphology and function of the ASER neuron and that axon placement defects of the ASER, observed in KNL-1 degrader animals could be attributed to a specific function of KNL-1 during axon development.

### Disruption of microtubule organization in the ASER neuron does not result in axon placement defects

To determine the underlying cause of ASER axon placement defects observed in KNL-1 degrader animals, we compared these defects with disruptions in cellular processes that affect neuronal morphology in *C. elegans*. We first examined the effect of disrupting microtubule organization, since the primary role of kinetochore proteins in dividing cells is to stabilize chromosome-microtubule connections during chromosome segregation (Cheerambathur and Desai, 2014). Previous studies had shown that mutations in Patronin/PTRN-1, the microtubule minus end stabilizer, led to morphological defects in *C. elegans* neurons (Marcette *et al*., 2014; Richardson *et al*., 2014). To understand its effects on axon placement, we analysed axonal trajectory in animals where *ptrn-1* was deleted. We found no significant difference in axon placement between the λ1*ptrn-1* and the control animals (**Figure 1C, D and S1A**). Similarly, deletion of NOCA-1, the orthologue of Ninein and minus end anchoring protein, crucial for microtubule organization in somatic cells of *C. elegans,* had no impact on axon placement of ASER (**Figure S1A,B and C**) (Wang *et al*., 2015). Additionally, we investigated the consequence of depleting the key microtubule nucleator γ-TuRC (γ-tubulin ring complex) to ASER axon structure and placement. γ-TuRC mediated nucleation of non-centrosomal microtubules is required to generate the microtubule arrays in axons and dendrites (Weiner *et al*., 2021). The depletion of core components of γ-TuRC such as GIP-2/GCP2 or GIP-1/GCP3 by ZIF-1 mediated protein degradation methods were previously shown to be sufficient to severely perturb microtubule organization in somatic tissues of *C. elegans* (Wang *et al*., 2015, 2017; Sallee *et al*., 2018). Degradation of endogenous GIP-2::GFP using the GFP nanobody::ZIF-1 based degron system resulted in a range of axonal defects in the ASER neuron, such as axon misrouting and ectopic branching (**Figure S1D and E**). However, the axon placement relative to the cell body and pharynx remained similar to control animals (**Figure 1C, D and S1A**). These findings collectively suggest that disruption of microtubule organization was not sufficient to explain the placement defects observed for KNL-1 degradation.

### Axon placement defects observed in KNL-1 degrader animals are similar to those seen with mutations in the cell adhesion protein, SAX-7/L1CAM

Next, we compared the axon placement defects in KNL-1 degrader animals to those caused by mutations in genes previously linked to guidance defects in amphid neurons (Zallen *et al*., 1999). Remarkably, our analysis revealed that the axon placement defects observed in KNL-1 degrader animals closely resembled those seen in animals with the deletion *nj48* allele of *sax-7* (Zallen *et al*., 1999; Sasakura *et al*., 2005). SAX-7 is a transmembrane protein composed of multiple Ig domains that facilitate homophilic interactions between neighbouring cells. In *C. elegans*, SAX-7 is crucial for axon guidance and positioning as well as the maintenance of the structural integrity of the nerve ring (Zallen *et al*., 1999; Sasakura *et al*., 2005; Bénard *et al*., 2012). Similar to KNL-1 degrader animals, in the *sax-7(nj48)* allele, the guidance of the ASER axon ventrally or its entry into the nerve ring was not affected. However, in both KNL-1 degrader animals and *sax-7(nj48)* mutants the axon was displaced more posteriorly in relation to the cell body and pharynx (**Figure 1C, D and S1A**). The intriguing similarity between the loss of function phenotypes between KNL-1 degrader animals and *sax-7(nj48)* mutants suggests that KNL-1 has distinct role in axon development and regulates axon guidance.

### Degradation of KNL-1 leads to a disorganized *C. elegans* nerve ring

Next, we explored the role of KNL-1 in organizing the axons within the nerve ring bundle. In our previous work, we had shown that endogenously tagged KNL-1::GFP was enriched in the developing dendrites in the embryonic stage (Cheerambathur *et al*., 2019). However, its expression in multiple neurons and the surrounding tissues had precluded us from observing specific axonal pools. Hence, to visualize KNL-1 selectively in the axons within the developing nerve ring, we used a split-GFP system (**Figure 2A**) (Kamiyama *et al*., 2016). The GFP1-10 was driven under an *hlh-16* promoter which is active in a subset of amphid and pioneer neurons that form the nerve ring. Consistent with our previous work, we observed GFP::KNL-1 in the dendrites of neurons in the embryonic stage (**Figure S2A and B**). Importantly, we detected the presence of KNL-1 in the developing axons in the embryos and within the axons of the nerve ring at the L1 stage (**Figure 2B, S2A and B**).

**Figure 2.**
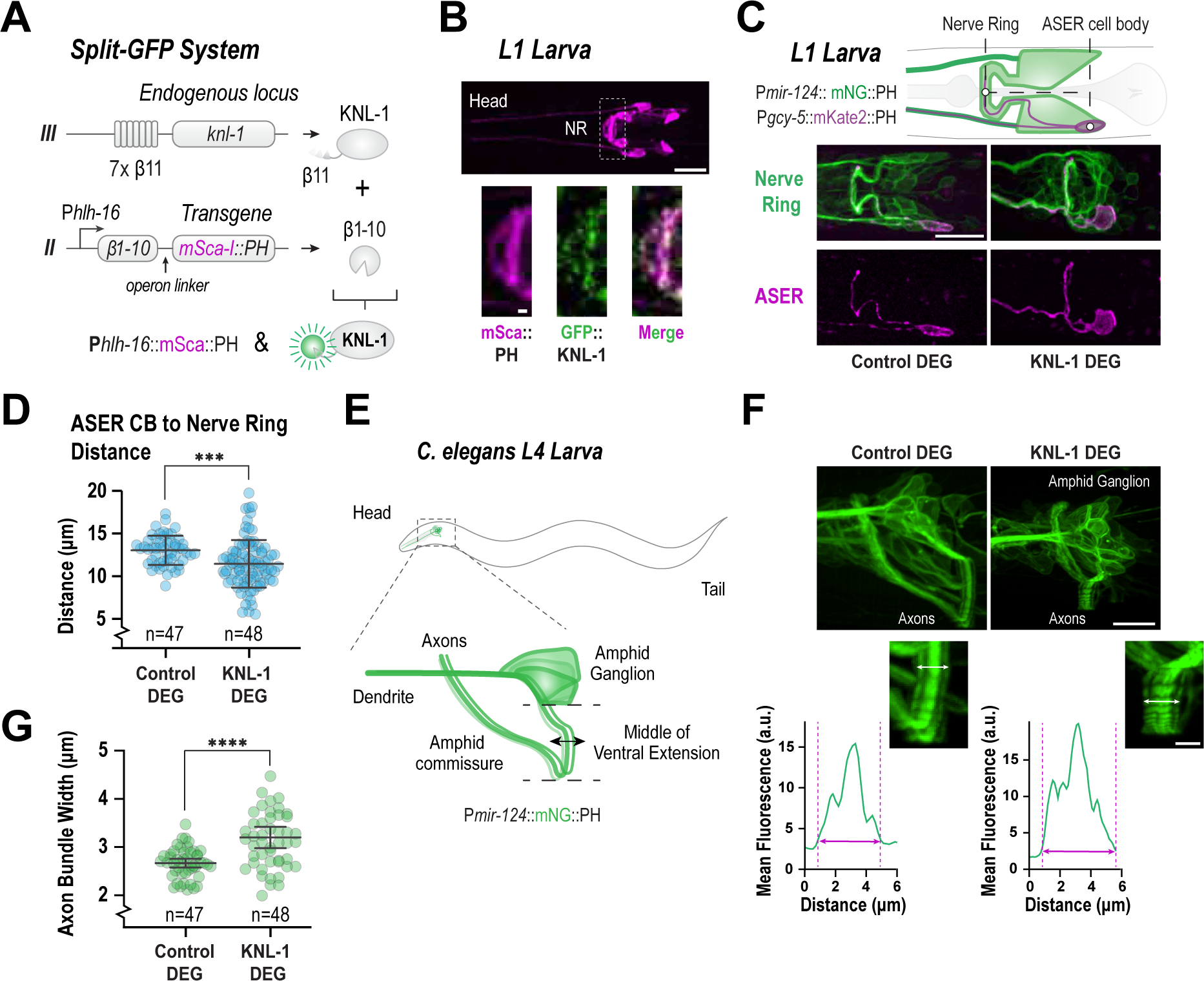
KNL-1 is required for proper architecture of the *C. elegans* nerve ring. A) Schematic of the split-GFP system used to express KNL-1 in a subset of head neurons. KNL-1 was endogenously tagged with seven repeats of GFP β11 at the N-terminus. β1-10 of GFP, linked to mScarlet-I::PH via the gpd-2/3 operon linker, was expressed under the *hlh-16* promoter. The *hlh-16* promoter is expressed in the amphid neuron, AWC, alongside other head neurons (Rapti et al. 2017). (B) Localization of GFP::KNL-1 in the neurons of an L1 stage animal using the *hlh-16* promoter driven split-GFP system. The upper panel displays the topology of the neurons expressing the membrane marker, mScarlet-I::PH. The below insets indicate the region of the nerve ring (NR) where GFP::KNL-1 is present. Scale bar, 10 μm (top), 1 μm (inset). (C) Schematic illustrating the positions of ASER (magenta) and the P*mir-124* expressing sensory neurons (green) (top). Images of the sensory neurons (green) and ASER (magenta) in the control and KNL-1 DEG conditions. P*mir-124* is expressed from embryogenesis until adulthood and is active in about 40 neurons, including the amphid neurons (Clark et al. 2010). In the control, the ASER axon, follows the same trajectory as the other sensory neurons. However, in KNL-1 DEG animals, the ASER axon remains within the nerve ring but is misplaced, relatively to its cell body. Scale bar, 10μm. (D) Quantification of the distance between the middle of the ASER cell body and the nerve ring. n indicates the number of animals. *** indicates p<0.001. (E) Schematic representation of the head sensory neurons expressing *Pmir-124*::mNeonGreen::PH transgene in an L4 larva, lateral view. (F) Images of the sensory neurons of control and KNL-1 DEG animals expressing the mNG::PH marker (top); the panels are corresponding zoomed views of the amphid commissure (middle). The white arrows show the measured distance in each bundle. The graphs below represent linescans of the fluorescence intensity of the corresponding commissure width. The pink arrows on the graph indicate the measured distance. Scale bar, 10 μm (top), 2.5 μm (crop-out). (G) Plot illustrating the thickness of the axonal commissure corresponding to the linescans in F. n represents the number of animals. Error bars denote 95% confidence interval. **** indicates p<0.0001.

Having observed an axon placement defect with the ASER neuron, we next investigated whether loss of KNL-1 affected the nerve ring structure and positioning. To do this, we engineered a strain expressing plasma membrane markers, under two different promoters, mNeonGreen::PH under *mir-124* promoter, active in all amphid neurons, and P*gcy-5*::mKate2::PH expressed only in the ASER neuron. This strain allowed us to track the position of the nerve ring axon bundle in relation to the cell body of the ASER neuron. In the control animals, the nerve ring was positioned ∼13 +/− 0.7μm anteriorly relatively to the ASER cell body at the L1 stage (**Figure 2C and D**). By contrast, in KNL-1 degrader animals, the nerve ring was displaced more posteriorly, ∼11.4 +/− 0.8μm away from the cell body. This positional change of the nerve ring was also observed in relationship to the pharynx bulb in the KNL-1 degrader animals as compared to the control, suggesting that KNL-1 function was required for the correct placement of the nerve ring (**Figure S2C**).

In addition to the misplaced nerve ring, the axons and cell bodies appeared to be disorganized in the KNL-1 degrader animals compared to the control (**Figure 2C)** in the L1 animals. Notably, the defects in nerve ring structure were variable across KNL-1 degrader animals making it challenging to characterize it quantitatively. To better define the axonal disorganization, we focused on the ventral extension of the amphid commissure, an axon rich tract, primarily formed by the axons projecting from the amphid neuron cell bodies. At the L4 stage, the amphid commissure was distinctly visible as a thick bundle in the strain expressing mNeonGreen::PH under the *mir-124* promoter (**Figure 2E, F and G**). The amphid commissure in the control animals showed a stereotypical organization where the axons project ventrally to form a tightly bundled structure of ∼2.5 µm in width (**Figure 2E, F and G**). In the KNL-1 degrader animals, although the amphid commissure extended ventrally, the axons within the commissure appeared to be dispersed. The width of the axon bundle in KNL-1 degrader animals was significantly higher than the control (**Figure 2F and G**). Taken together, these results suggest that KNL-1 plays a key role in facilitating proper axonal bundling and ensures the correct placement of the nerve ring during development.

### Neuronal function of KNL-1 relies on the signalling and microtubule binding modalities essential to chromosome segregation

To address the mechanism by which KNL-1 facilitates nerve ring assembly, we adopted a structure-function approach to identify the functional modules within KNL-1 important for its neuronal function. In mitosis, the N-terminal half of the KNL-1 serves as a docking site for protein phosphatase 1 (PP1), and the spindle assembly checkpoint protein, BUB-3/BUB-1, while the C-terminus of KNL-1 recruits the Ndc80 complex (**Figure 3A**) (Espeut *et al*., 2012, 2015; Moyle *et al*., 2014).

**Figure 3.**
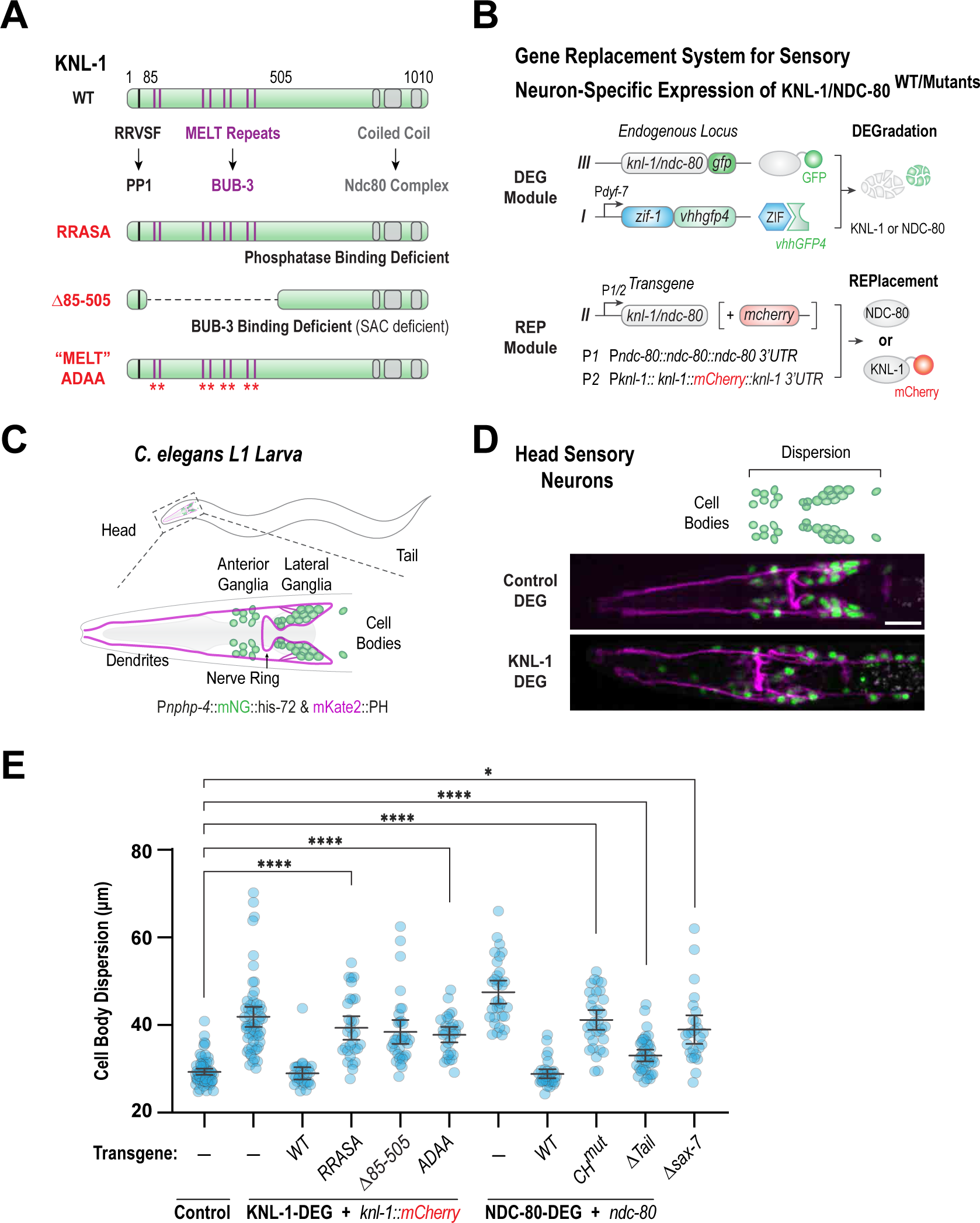
Neuronal function of KNL-1 requires both the microtubule binding and signaling activities. (A) Schematic illustrating the structure of KNL-1 (top) and the three mutants of KNL-1 used in the study: RRASA, Δ85-505 and ‘’MELT‘’ ADAA (bottom). (B) Gene replacement system used to assess the effect of various domains of KNL-1 and NDC-80 on the structure of head sensory nervous system. The P*dyf-7* driven tissue-specific degradation system was used to selectively degrade endogenously GFP-tagged KNL-1 or NDC-80 in the head sensory neurons. The degrader-resistant *knl-1*::mCherry or *ndc-80* wild type and mutant transgenes were expressed under endogenous promoters. (C) Schematic of the ciliated neurons at the L1 larval stage. To visualize the cell bodies (green) and the membranes (magenta) of these neurons, an operon construct that expressed mNG::his-72 and mKate2::PH under the control of *nphp-4* promoter was used. The cell bodies of the ciliated neurons are organized into anterior and lateral ganglia, while their axons form part of the nerve ring. (D) Images of head sensory neuron nuclei (green) and plasma membranes (magenta) for the indicated conditions. Scale bar,10 μm. (E) Quantification of dispersion of sensory neuron cell bodies in the indicated conditions. n represents the number of animals. Error bars denote 95% confidence interval. * and **** indicates p<0.05 and 0.0001, respectively.

To investigate the importance of the signalling modules within KNL-1 in the neurons, we developed a gene replacement system that allowed us to selectively express “separation of function” alleles of KNL-1 with compromised activities in sensory neurons (**Figure 3A and B**). Previous studies in mitosis have demonstrated that the mutation of PP1 docking site RRVSF motif to RRASA abrogates PP1 recruitment, while deletion of the region within KNL-1 containing the MELT repeats (τι85-505), or mutations of 9 MELT repeats to ADAA diminish BUB-3 binding to mitotic kinetochores and abolish the spindle assembly checkpoint (**Figure 3A**)(Espeut *et al*., 2012; Moyle *et al*., 2014). The degrader-resistant KNL-1 alleles were expressed as single copy transgenes under endogenous *knl-1* promoter, and we confirmed the presence of transgenes in the embryonic head neurons using fluorescence microscopy (**Figure S3A**). Next, to assess the role KNL-1 signalling modules in neuronal development, we utilized a previously established fluorescence-based assay to evaluate the structural integrity of head sensory neurons during the L1 larval stage (Cheerambathur *et al*., 2019). We labelled the nuclei and plasma membranes of a select group of ciliated sensory neurons, which included amphid neurons, by employing mNeonGreen::Histone and mKate::PH markers under the control of the *nphp-4* promoter (**Figure 3C**). By the L1 stage, in control animals, the cell bodies within the anterior and lateral ganglia were tightly clustered in an ∼30 µm wide region on either side of a tightly bundled nerve ring (**Figure 3D and E**). Consistent with our previous findings, this stereotypical architecture was perturbed after the degradation of KNL-1. When we expressed the KNL-1 PP1 binding mutant (RRASA) or BUB-3 binding/spindle assembly checkpoint deficient mutants (ϕλ85-505 and ADAA), we observed significant dispersion of cell bodies of the ganglia towards the anterior and posterior directions similar to the degradation of KNL-1 (**Figure 3E**). These experiments strongly suggest that the RVSF and MELT repeat motifs on KNL-1 are critical to its neuronal function and that the ability of KNL-1 to recruit signalling proteins appears to be important for its proper function in the nervous system.

The Ndc80 complex, the core microtubule coupler at the mitotic kinetochore binds to the C-terminus of KNl-1. Similar to KNL-1, NDC-80::GFP was present in the axons during nerve ring assembly (**Figure S2B**). As shown previously, degradation of NDC-80 subunit or mutant transgenes compromised in microtubule binding, fully recapitulated the loss of KNL-1 and the KNL-1 signalling domain mutant alleles in head sensory nervous system architecture (**Figure 3E**). Although technical challenges precluded us from directly assessing whether the recruitment dependencies observed in mitotic kinetochore are valid in neurons, the phenotypic similarity of the KNL-1 and NDC-80 degraders suggest that neuronal function is dependent on the same interface of the mitotic kinetochore responsible for chromosome-microtubule attachment (**Figure 3A).** More importantly, there appears to be a crosstalk between the signalling and microtubule binding domains within the KMN network in the neurons.

The dispersal of neuronal cell bodies within the ganglia following the loss of KNL-1 and NDC-80 indicates that the kinetochore proteins might have roles in regulating cell-cell contacts. To understand the mechanism by which KNL-1 and Ndc80 complex influences the neuronal cell body organization, we assessed the cell body dispersion defect seen with loss of kinetochore proteins to that of *sax-7(nj48)* mutant. Importantly, previous studies have implicated SAX-7 in head ganglia organization (Sasakura *et al*., 2005; Bénard and Hobert, 2009) and the axon placement defects were strikingly similar between the *sax-7(nj48)* allele and KNL-1 and NDC-80 depletion. While the cell body organization was disrupted in the *sax-7(nj48)* mutant at the L1 stage, the severity of the defect was significantly less than that observed in KNL-1 or NDC-80 degraded animals (**Figure 3E and S3C**). Prior studies have shown that *sax-7* defects are progressive and more pronounced at later stages of development and that additional molecular factors such as extracellular matrix proteins influences the head ganglia structure (Bénard *et al*., 2006, 2012). Therefore, the ganglia disorganization seen in the absence of KNL-1 and NDC-80 may be independent of SAX-7. Nevertheless, these findings imply that the neuronal function of kinetochore proteins may be cell nonautonomous and that they operate within a multicellular framework to shape the architecture of the nervous system.

### Microtubule organization of the ASER neuron is unaffected in KNL-1 degrader animals

We next investigated whether the axonal function of KNL-1 might be associated with regulating microtubule organization and dynamics in the axons. We had previously analysed the microtubule plus end dynamics in the absence of KNL-1 by tracking the microtubule end-binding protein EBP-2 (EB1) in the amphid dendrite cluster during embryogenesis (Cheerambathur *et al*., 2019). However, axons differ in their microtubule organization and their microtubule features were not examined in this prior study. Furthermore, due to the presence of a microtubule organizing centre at the ciliary base of the amphid dendrites, the previously observed EBP-2 dynamics in the developing dendrites predominantly reflects microtubule nucleation occurring from the ciliary base (Garbrecht *et al*., 2021; Magescas *et al*., 2021). To monitor axonal microtubule orientation and dynamics we expressed EBP-2 fused to mNeonGreen in ASER amphid sensory neuron and tracked the movement of EBP-2::mNG puncta at the L1 stage (**Figure 4A** and **S4A**). In the amphid neurons, microtubule plus ends in the axons are directed away from the cell body whereas in the dendrite, microtubule plus ends are oriented towards the cell body (Maniar *et al*., 2012; Harterink *et al*., 2018). In control animals, all axonal EBP-2::mNG puncta moved away from neuronal cell bodies, whereas all dendritic puncta moved towards the cell bodies (**Figure 4B** and **S4B**). This pattern of EBP-2 movement remained unaltered in KNL-1 degrader animals suggesting that microtubule orientation in the neuron is unaffected in the absence of KNL-1. To further validate our findings, we examined the distribution of presynaptic axonal protein mKate2::SNB-1 in the ASER neuron at the L4 larval stage (**Figure 4E**). Presynaptic proteins such as SNB-1 are localized exclusive to axons and disruption of neuronal microtubule polarity results in the redistribution of axonal proteins to dendrites (Nonet, 1999; Maniar *et al*., 2012). We found that SNB-1 distribution was similar in both control and KNL-1 degrader animals, with the majority of mKate2::SNB-1 signal confined to the axons of ASER neuron (**Figure 4F and G**). As a control, we analysed SNB-1 distribution in the ASER neuron after degradation of GIP-2::GFP which has previously been shown to impact microtubule dynamics within the dendrites (Harterink *et al*., 2018). After GIP-2 degradation, significant percent of neurons, exhibited mislocalization of SNB-1 in the dendrites of the ASER neuron (**Figure S4E and F**). Next, we measured the microtubule dynamics of the growing plus ends in the ASER axon by tracking EBP-2 comets. Specifically, we observed no change in the frequency or velocity of the EBP-2 comets, suggesting that there is no change in the dynamics of growing plus ends of microtubules (**Figure 4C and D**). Thus, these observations suggest that KNL-1 degradation in the sensory neurons does not alter microtubule polarity and dynamics and that KNL-1’s function in the neurons is not associated with a role in regulating the overall stability or organization of microtubules.

**Figure 4.**
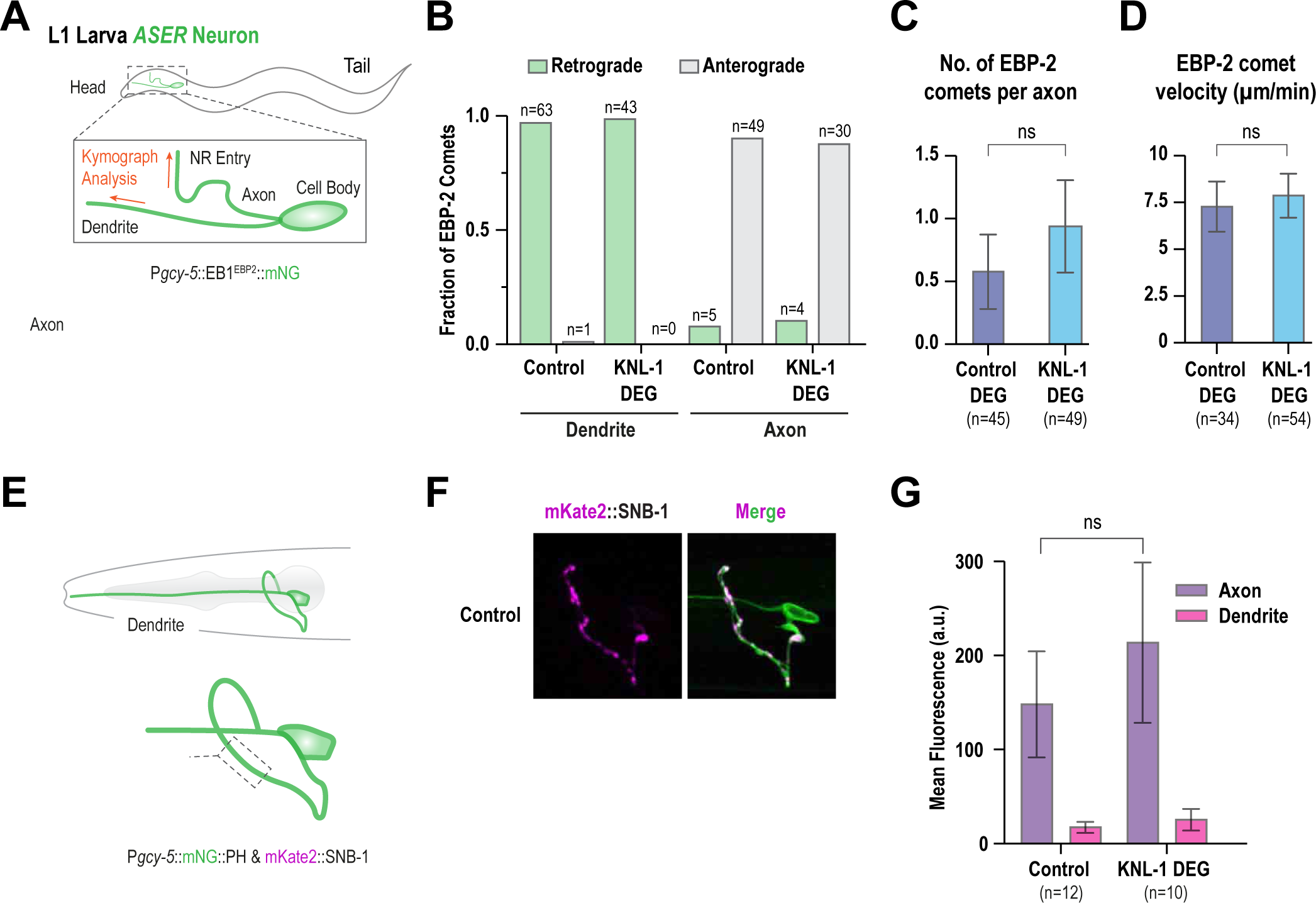
Microtubule organization within the neuron is intact after KNL-1 degradation. (A) Cartoon shows the polarity of microtubules in the axon and dendrite of the ASER neuron at the L1 larval stage. Microtubule plus ends in the ASER neuron were labelled using EB1^EBP-2^::mNG expressed under the *gcy-5* promoter. Orange arrows indicate the direction of the kymographs, in the dendrite and axon. (B) Quantification of EB1^EBP-2^ comet direction within the dendrite and axon of the control and KNL-1 DEG animals. (C-D) Quantification of comet number and velocity in the axon in the indicated conditions. n represents number of axons. ns indicates non-significant. (E) Schematic showing the localization of synaptic marker SNB-1 in the ASER axon. To visualize ASER neuron morphology and SNB-1 protein localization, the transgenes P*gcy-5*::mNeonGreen and P*gcy-5*::mKate2::SNB-1 were utilized, respectively. SNB-1 is a protein involved in synaptic vesicle-membrane fusion and is typically found in puncta along the axon of presynaptic neurons. (F) Localization of synaptic marker SNB-1 (magenta) in the axon of the ASER (green) in control and KNL-1 DEG appears similar. Scale bar,10 μm. (G) Quantification of SNB-1 signal in the axon, in the indicated conditions. Error bars denote 95% confidence interval. n represents the number of animals. ns indicates non-significant.

### NDC-80 microtubule binding domain facilitate axon bundling during early stages of nerve ring assembly

Since the primary function of KNL-1 and Ndc80 complex appears to be associated with the bundling of axons in the nerve ring, we focused on understanding the source of underlying structural defects. The nerve ring assembly in *C. elegans* embryos initiates during bean stage and culminates towards the end of embryogenesis (Rapti *et al*., 2017; Moyle *et al*., 2021). A key step in nerve ring formation involves the extension of axonal outgrowths by a group of 8 pairs of bilaterally symmetrical pioneer neurons to generate a ring like scaffold (**Figure 5A**). Subsequent to this, other neurons, including the sensory neurons like the amphids, project their axons in a hierarchical fashion to generate a functionally stratified nerve ring (**Figure S5A**). To explore the involvement of KNL-1 and NDC-80 in this process, we performed cell-specific labelling of both proteins in the pioneer neurons using the split-GFP system. We found that both KNL-1 and NDC-80 were enriched in the pioneer axons during nerve ring assembly (**Figure 5B**). This led us to hypothesize that KNL-1 and NDC-80 might play a role in bundling the axons, that build the nerve ring scaffold. To test this, we attempted to create a pioneer neuron-specific GFP nanobody::ZIF-1 fusion based protein degradation system. Unfortunately, we encountered difficulties in identifying a promoter that would express the GFP degrader at the appropriate time before ring assembly. As an alternative approach, we created partial loss of function alleles of NDC-80 and KNL-1, that compromised specific functions but generated viable animals that allowed us to track nervous system development. We particularly focused on the microtubule binding domains of the NDC-80 given its importance for neuronal function. The microtubule binding activity of the Ndc80 complex primarily resides in the unstructured N-terminal tail region of the NDC-80 subunit, known as the NDC-80 Tail, and the calponin homology (CH) domain of the NDC-80 subunit (**Figure 5C**) (Ciferri *et al*., 2008; Alushin *et al*., 2010). Both microtubule domains within the Ndc80 complex are critical for its neuron function (Cheerambathur *et al*., 2019)). We succeeded in generating a viable animal lacking the basic N-terminal tail of NDC-80 which is essential for Ndc80 complex binding to microtubules *in vitro*. Although, the N-terminal tail has previously been shown to contribute to regulation of chromosome microtubule attachments, deletion of the NDC-80 Tail did not seem to have any discernible effect on chromosome segregation (**Figure S5B**) (Cheerambathur *et al*., 2013, 2017).

**Figure 5.**
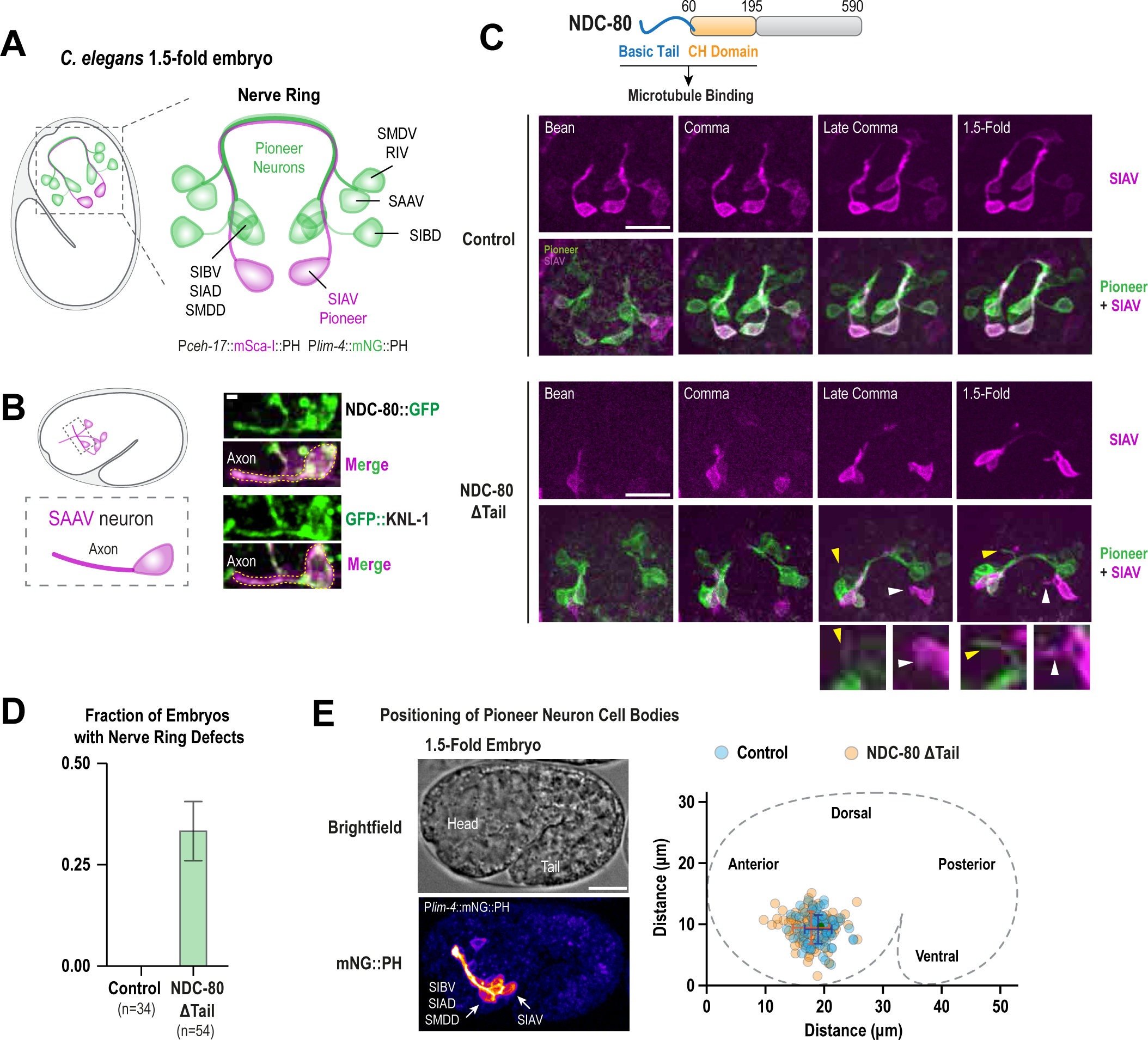
NDC-80 microtubule binding domain is important for axonal bundling during early stages of nerve ring assembly. (A) Schematic shows the lateral view of a *C. elegans* embryo at 1.5-fold stage and that of the pioneer neurons that form the scaffold of the nerve ring. Pioneer neurons were labelled using P*lim-4*::mNG::PH, which is expressed in 8 pairs of pioneer neurons (SIBV, SIAD, SIAV, SIBV, SAAV, SMDD, SMDV and RIV) during embryogenesis. P*ceh-17*::mSca-I::PH expression was restricted to the SIAV pioneer neuron. (B) Schematic showing the *hlh-16* expressing neurons in the embryo (left). P*hlh-16* is expressed in the pioneer neuron SAAV. Localization of NDC-80::GFP and GFP::KNL-1 (green) in the SAAV neuron using the *hlh-16* promoter driven split-GFP system. The membranes of the neurons are shown in magenta using the membrane marker, mScarlet-I::PH. Scale bar, 1 μm. (C) Domain organization of NDC-80 (Top). (Below)Time-lapse sequences showing the progression of axon extension in pioneer neurons from bean stage to the 1.5-fold stage of embryonic development, in control and NDC-80 ΔTail. The SIAV neuron (magenta) extends its axon later, compared to the other pioneer neurons. It eventually enters the pioneer axon bundle (green) which has already been formed by the axons of SIBV, SIAD, SIBV, SAAV, SMDD and SMDV. Notably, the expression of the *lim-4* promoter driven marker in the SIAV neuron appears faint at this stage compared to the marker expressed under *ceh-17* promoter. The process of ring formation is completed around the 1.5-fold stage. In NDC-80 ΔTail embryos, some axons fail to incorporate into the bundle and the SIAV axon frequently fails to enter the nerve ring. The insets below the panel indicates the failure of non-SIAV pioneer axon incorporation (yellow arrowhead) and the failure of the SIAV axon to extend and enter the nerve ring (white arrowhead). Scale bar, 10 μm. (D) Quantification of the embryos that present axon bundling defects. n represents the number of embryos. Error bars depict the standard error of the proportion. (E) Plot illustrates the positioning of the cell bodies of pioneer neurons SIBV, SIAD, SMDD and SIAV in the embryo. Images of the brightfield of a 1.5-fold embryo (left top) and the pioneer neurons labelled by P*lim-4*::mNG::PH (left bottom). The graph (right) illustrates the X,Y coordinates of each individual neuronal cell body in the embryo. n represents the number of cell bodies. The error bars denote the 95% confidence interval. Scale bar, 10 μm.

Next, we monitored axon assembly during nerve ring formation in wild type and NDC-80 ΔTail animals. We developed a strain that enabled us to track axon extension in all pioneer neurons (P*lim-4*::mNeonGreen::PH) as well as specifically the incorporation of the SIAV pioneer axon (P*ceh-17*::mScarlet-I::PH) into the ring scaffold and used time-lapse imaging to characterize the ring assembly with high temporal resolution. In both the wild type and the NDC-80 ΔTail embryos, the pioneer neurons extended their axonal growths towards the anterior of the embryo at the bean stage, eventually forming a ring like scaffold by 1.5-fold (**Figure 5C**). However, in NDC-80 ΔTail embryos we noticed several issues with axon incorporation and bundling (**Figure 5C and D**). In a significant proportion of NDC-80 ΔTail embryos, axons of some neurons failed to incorporate correctly or were splayed from the bundle (yellow arrowheads in **Figure 5C**). Additionally, using our dual-colour marker that enabled us to visualize the trajectory of the individual SIAV axon specifically into the pioneer axon scaffold (white arrowheads in **Figure 5C and S5C**) we observed the SIAV growth cones either failing to extend or being correctly guided towards the pioneer neuron cell body cluster, but its axon stopping the extension and failing to integrate in the axonal bundle (**Figure 5C** and **S5C**). Considering our earlier observations that the cell body organization in sensory neuron ganglia in KNL-1 and NDC-80 degrader animals is impaired, we also monitored the positioning of four pioneer neuron cell bodies (SIBV, SIAD, SMDD and SIAV) in wild type and NDC-80 ΔTail embryos (**Figure 5E**). Our analysis revealed no difference in the organization between the wild type and NDC-80 ΔTail embryos. In both cases the cell bodies clustered at the ventral side of the embryo head and showed no significant difference in their distribution. These results suggest that the microtubule binding domain of NDC-80 actively regulates the axon bundling during nerve ring assembly, and that the Ndc80 complex plays a crucial role in building a proper nerve ring scaffold.

## DISCUSSION

The wiring of the nervous system is a complex process with significant implications towards animal behaviour, yet its molecular underpinnings remain poorly understood. Here we highlight a non-canonical function for the conserved KMN network components, the KNL-1 and the Ndc80 complex, in the formation and patterning of the head sensory nervous system in *C. elegans*. Specifically, the kinetochore proteins are required for precise positioning and bundling of the axons within the *C. elegans* nerve ring and for the maintenance of neuronal cell body clusters within the sensory ganglia. Direct phenotypic comparisons revealed the axonal function of the KNL-1 and Ndc80 complex is likely mechanistically distinct from that of microtubule array organizers in neurons, such as Patronin or the γ-tubulin ring complex. In our previous work, we found that degradation of KNL-3, a component of the Mis12 complex, also disrupted the organization the nerve ring ganglia akin to the effects seen with depletions of KNL-1 and NDC-80. Taken together, these observations suggest that the entire KMN network may act at a multicellular level to organize and maintain the *C. elegans* head sensory nervous system (Cheerambathur *et al*., 2019). This atypical role for KMN network in facilitating the tissue architecture of the central nervous system seems universal, as evidenced by similar organizational defects in *Drosophila* brain structure with knockdown of the Mis12 complex (Zhao *et al*., 2019).

The role of KMN proteins in higher-order neuronal organization is further supported by the striking resemblance between the phenotypes resulting from KNL-1 depletion and the loss of SAX-7, a cell adhesion molecule critical for neuronal tissue adhesion. Loss of cell adhesion (CAM) or extracellular matrix (ECM) family proteins in *C. elegans* also results in mispositioning of neuronal cell bodies from the ganglia and/or misplacement of axons within the axonal tracts (Aurelio *et al*., 2002; Sasakura *et al*., 2005; Bénard *et al*., 2012; Kim and Emmons, 2017). The structural defects due to loss of CAM and ECM proteins were primarily observed in older animals and has been attributed to an inability to counteract mechanical forces arising from locomotion. Nevertheless, the phenotypic similarity with the kinetochore is indicative of a role for KMN proteins in coupling adjacent cells within the neuronal tissue, likely by influencing the microtubule cytoskeleton within the neurons. Interestingly, SAX-7 has been found to directly link with cytoskeletal proteins such as UNC-44 via the ankyrin domain to promote the structural organization of head ganglia in *C. elegans* suggesting that the connection between CAMs and the cytoskeleton is key to preserve the organization of nervous system (Pocock *et al*., 2008; Zhou *et al*., 2008).

While the precise mechanism by which the microtubule cytoskeleton facilitates neuronal adhesion is not well understood, microtubules have emerged as key regulators of cell adhesion in other cellular contexts (Seetharaman and Etienne-Manneville, 2019). For example, in cell migration, microtubules regulate the turnover of focal adhesions through interactions with integrin-mediated adhesion and a feedback between cell-cell adherens junctions and microtubule dynamics exists in newt lung epithelial cells (Waterman-Storer *et al*., 2000; Stehbens and Wittmann, 2012; Seetharaman *et al*., 2022). Given that KMN proteins bind to and stabilize chromosome-microtubule attachments at the mitotic kinetochore, it is plausible they could coordinate the link between neuronal cell coupling and the microtubule cytoskeleton by stabilizing microtubule contacts at the cell cortex. Although we did not observe changes to overall microtubule orientation or dynamics within the axons of L1 animals after degradation of KNL-1, KMN proteins may regulate microtubule dynamics in a specific axonal region during earlier developmental stages not captured by our analysis. Supporting this, the failure of the NDC-80 microtubule binding mutant NDC-80 ι1Tail to bundle axons implies that interaction between Ndc80 complex and microtubules is involved in modulating axon adhesion in the developing nerve ring.

Apart from the chemical signals that guide axon growth and elongation, recent studies have implied that mechanical signals also contribute to this process. The axonal growth of the retinal ganglion cells in *Xenopus laevis* is shown to depend on the mechanical environment of the neurons and mechanical tension shapes axonal fasciculation in mouse neural explants (Koser *et al*., 2016; Šmít *et al*., 2017). Furthermore, the mechanical tension sensing in axons appears to involve interactions between cell adhesion molecules and the cytoskeleton, analogous to that proposed for growth cone motility (Suter and Forscher, 1998). Notably, kinetochores are well-known tension sensors that detect pulling forces exerted by spindle microtubules on sister chromatids during chromosome segregation (Sarangapani and Asbury, 2014; Lampson and Grishchuk, 2017). *In vitro* experiments using reconstituted mitotic kinetochore particles have shown that tension can modulate kinetochore-microtubule attachment stability similar to the “catch bond” mechanism observed with some receptor-ligand protein pairs that mediate cell adhesion (Marshall *et al*., 2003; Akiyoshi *et al*., 2010). The catch-bond like behaviour exhibited by kinetochore proteins *in vitro* is attributed, in part, to the ability of microtubule binding proteins, like the Ndc80 complex, to interact and modulate microtubule plus end dynamics (Sarangapani and Asbury, 2014). The modulation of microtubule dynamics is known to influence axonal outgrowth (Voelzmann *et al*., 2016; de Rooij *et al*., 2018; Alfadil and Bradke, 2023) and a feedback loop between mechanical tension and microtubule dynamics plays a central role in modulating the adhesive properties of cells at focal adhesions (Seetharaman and Etienne-Manneville, 2019). While we do not know the exact functional and physical relationship between cell adhesion molecules and microtubules in the axonal context, it is intriguing to think that crosstalk mediated by KMN proteins could contribute to the tension-dependent monitoring mechanism during nerve ring axon bundle formation. For instance, changes in tension detected by KMN proteins may alter cytoskeletal dynamics (e.g., microtubule polymerization), which then could influence CAMs and ECM proteins to weaken or strengthen axon-axon contacts. In order to test the above hypotheses, it would be important to identify the neuronal interactors of KMN proteins.

The domain analysis of KNL-1 revealed that the KNL-1 N-terminus known for overseeing the fidelity of chromosome-microtubule attachments also plays an important role in maintaining the head sensory nervous system architecture. Specifically, two key domains – the docking motif of protein phosphatase 1 (PP1), and the MELT repeats that activate the spindle assembly checkpoint (SAC) pathway – are essential for the neuronal function of KNL-1. During mitosis, the SAC monitors microtubule occupancy at the kinetochore and coordinates the integrity of chromosome-microtubule attachments with cell cycle progression (Lara-Gonzalez *et al*., 2021). In the absence of microtubule attachments, phosphorylation of KNL-1 MELT repeats by the mitotic kinases (Mps1/Plk1) recruits the BUB proteins to catalyse the formation of the Mitotic Checkpoint Complex (MCC) (London *et al*., 2012; Shepperd *et al*., 2012; Yamagishi *et al*., 2012). The MCC composed of Mad2, Cdc20, BubR1 and Bub3 serves as an inhibitor of Anaphase Promoting Complex (APC), an E3 ubiquitin ligase that promotes mitotic exit. Upon microtubule attachment, the SAC is silenced through various mechanisms including the localization of PP1 to the KNL-1. The PP1 recruitment correlates with the state of kinetochore-microtubule attachment and is proposed to dephosphorylate the MELT repeats thereby stopping SAC signalling (Liu *et al*., 2010; Meadows *et al*., 2011; Roy *et al*., 2019). The importance of KNL-1 modules used for SAC activation and silencing pathways suggests that effective communication and coordination between the microtubule attachment and signalling machinery is also key to neuronal development. Interestingly, the signalling arm of the kinetochore (including SAC) has been observed to play a post-mitotic role in neuronal context in other species. For example, in *Drosophila* larval neurons both KMN and SAC proteins are required to promote dendrite regeneration and PP1 phosphatase mediated regulation controls axon guidance in the hippocampal neuronal cultures (Hoffman *et al*., 2017; Hertzler *et al*., 2020; Kastian *et al*., 2023). Depletion of both KMN and SAC proteins in *Drosophila* neurons alters microtubule dynamics, indicating their involvement in detecting defective microtubule connections and neuron growth behaviour (Hertzler *et al*., 2020). The SAC signalling pathway in the neurons likely involves regulation of APC which in turn could regulate the turnover of target proteins that control nervous system structure. Support for this hypothesis comes from studies of post-mitotic roles for Cdc20/APC in neurons across organisms (Huang and Bonni, 2016). Cdc20/APC is required during dendrite morphogenesis of mouse cerebellar and hippocampal neurons and APC has been shown to regulate synaptic transmission in the neuromuscular junctions of *C. elegans* (Kim *et al*., 2009; Kowalski *et al*., 2014; Watanabe *et al*., 2014). Moreover, BUBR1 in post-mitotic mouse neurons has been found to modulate Cdc20/APC, with one of the direct targets of this interaction being FEZ1 (fasciculation and elongation protein zeta1), a protein involved in axon outgrowth and fasciculation (Watanabe *et al*., 2014). To understand how KMN signalling activity and microtubule coupling are integrated in the central nervous system, it would be critical to understand how SAC signalling reports microtubule states within neurons and whether it controls Cdc20/APC activity, and its putative neuronal targets. In conclusion, our study highlights the intricate and dynamic interplay between KMN proteins, microtubules, signalling and cell adhesion in shaping the neuronal architecture and further investigation is required to uncover the precise mechanisms involved.

## ACKNOWLEDGMENTS

We thank Ikue Mori for the *sax-7* allele; Kevin Hardwick, Rebecca Green and Pablo Lara-Gonzalez for comments on the manuscript; David Kelly and Toni McHugh (Wellcome Centre Optical Instrumentation Laboratory, (COIL)) for microscopy and image analysis support. Some strains were provided by the CGC, which is funded by NIH Office of Research Infrastructure Programs (P40 OD010440). This work was supported by a Sir Henry Dale Fellowship to D.K.C. jointly funded by the Wellcome Trust and the Royal Society (208833/Z/17/Z), a Sir Henry Wellcome Postdoctoral Fellowship (215925) awarded to B.P. and Darwin Trust PhD Studentship, Edinburgh awarded to V.R.O. COIL is supported by a Core Grant (203149) to the Wellcome Centre for Cell Biology at the University of Edinburgh.

## MATERIALS AND METHODS

### C. elegans strains

The *C. elegans* strains were grown on Nematode Growth Media (NGM) plates seeded with *E. coli* OP50 bacteria at 20°C. All strains used in the study are listed in Table S1.

### *C. elegans* transgenic strain construction

The Mos1 mediated Single Copy Insertion MosSCI method (Frøkjaer-Jensen *et al*., 2008) was used to generate transgenic animals stably expressing fluorescent markers, vhhgfp4::zif-1 degrader, *knl-1* and *ndc-80* transgenes. The transgenes were either cloned into pCFJ151 for insertion on chromosome I (*oxTi185*), II (*ttTi5605*), IV (*oxTi177*) or V (*oxTi365*). Information on the promoters used for expression of the transgenes in specific neurons or gene replacement of *knl-1* and *ndc-80* can be found in Table S2. Briefly, a mix of plasmids that contains the transgene of interest including a positive selection marker, a transposase plasmid, three fluorescent markers for negative selection [pCFJ90 (Pmyo-2::mCherry), pCFJ104 (Pmyo-3::mCherry) and pGH8 (Prab-3::mCherry)] were injected into *unc-119* mutant animals. Moving, non-fluorescent worms were then selected, and insertions were confirmed by PCR using primers spanning both homology arms.

The alleles *7Xgfp11::knl-1*, *AID::dark-gfp::knl-1* and *ndc-80(ΔTail)* were generated by CRISPR/Cas9-mediated genome editing using the Cas9/RNP method as described in (Paix *et al*., 2015). Table S3 lists the genomic sequences targeted by the guide RNAs. To generate *7Xgfp11::knl-1* and *AID::dark-gfp::knl-1* alleles, a double-stranded DNA repair template containing the desired insertion and 35bp homology arms template was custom-synthesized as a gBlock by Integrated DNA Technologies. For engineering the ndc-80(ΔTail) allele a single-stranded repair template with 35bp homology arm was used. Gene editing was performed by injecting the repair template and a ribonucleoprotein (RNP) mixture comprising crRNA/tracrRNA (Integrated DNA Technologies) and purified recombinant Cas9 enzyme into the gonads of young N2 adults. The injection mix also contained RNPs targeting the R92C mutation in dpy-10, which results in a dominant roller phenotype (Arribere et al., 2014), to select for animals with potentially successful edits. The roller worms were further screened by genotyping PCRs using primers that track the desired genome insertion (*7Xgfp11::knl-1*, *AID::dark-gfp::knl-1*) or deletion (*ndc-80(ΔTail*)). Positive clones from the PCR were then confirmed by sequencing.

### Fluorescence microscopy and analysis

For all imaging experiments, animals (L1 or L4 stage) were anaesthetized in 5mM levamisole, and both the larvae and embryos were mounted in M9 on 2% agarose pads. The imaging was done with a spinning disc confocal microscopy system equipped with a Yokogawa spinning disk unit (CSU-W1), a Nikon Ti2-E fully motorized inverted microscope and a Photometrics Prime 95B camera, unless stated otherwise. A CFI60 Plan Apochromat lambda 60X (Nikon) objective was used to image the neurons in the hatched animals whereas CFI60 Plan Apochromat lambda 100X Oil (Nikon) objective was used to image embryonic neurons. All images were processed using ImageJ (Fiji) software.

To image the ASER neuron morphology at the L4 stage, Z-stacks of 1.0 μm spacing were captured to cover the entire neuron expressing the P*gcy-5*::mKate2::PH marker (Figure 1, S1). Z-stacks were then projected into a maximum intensity projection (MIP) in ImageJ (Fiji) and the distance from the middle of the cell body or from the centre of the terminal pharynx bulb to the dendrite-axon cross-section point was calculated to measure the distance between the axon entry point to the nerve ring and the cell body of the ASER neuron.

The structure of sensory head nervous system in (Figure 2, 3), was imaged using L1 animals expressing the markers that labelled the membrane (mKate2::PH) and nucleus (mNeonGreen::his-72) of the ciliated neurons (P*nhp-4* promoter).Z stacks of size 25×0.5μm were acquired, and maximum intensity projections of the z-stacks were made in ImageJ (Fiji). To analyze the cell body within the head region, the distance between the most anterior and most posterior nucleus was calculated.

To image the axon organization within the amphid commissures (Figure 2), z-stacks spanning 29×0.75μm in the head region of L4 stage animals expressing P*mir-124*::mNeonGreen::PH were acquired. The width of the axonal bundle was determined by performing line scan analysis (line 1μm width) across the middle of the ventrally extended axonal amphid commissure using the maximum intensity projections of the neurons. The base width of this line scan was then used to calculate the width of the axonal bundle.

To image the localization of the various KNL-1 mutants fused to mCherry expressed in the background of the KNL-1 GFP degrader, z-stacks of size 21×0.5μm of 1.5-fold embryos were acquired. To quantify the signal of each KNL-1 mutant, the z-stacks were projected into maximum intensity projections and the mean fluorescence intensity (across the head of the embryo, 15μm radius circle) was then measured and plotted, using a ROI of 20×15μm.

The distribution of presynaptic marker, SNB-1 in the ASER neurons, (Figure 4), was assessed by collecting z-stacks of size 29×0.75μm of ASER neuron in L4 animals expressing mNeonGreen-PH and mKate2-SNB-1 under the *gcy-5* promoter. To quantify SNB-1 signal, the z-stacks were projected into maximum intensity projections and the mean fluorescence intensity was then measured and plotted along the length of the axon and 50um proximal to the dendrite, using a segmented line with a width of 3 pixels in Image J (Fiji).

For imaging the GFP::KNL-1 expression in the nerve ring (Figure 2), 15×0.5μm z-stacks of neurons expressing the mScarlet-I::PH marker L1 animals were acquired.

### Time lapse imaging & analysis

Time lapse images were acquired using a spinning disc confocal imaging system with a Yokogawa spinning disk unit (CSU-W1), a Nikon Ti2-E fully motorized inverted microscope, and Photometrics Prime 95B camera.

EBP-2 dynamics: For imaging the dynamics of mNeonGreen::EBP-2 of the ASER axon and dendrite, L1 animals were anaesthetized in 5mM levamisole and mounted in M9 on 2% agarose pads. Movies of 7×0.75μm z-slices of the ASER axon and dendrite were acquired at 1 frame per second using a CFI60 Plan Apochromat lambda 100X Oil (Nikon) objective (Figure 4 & S4). Kymographs of EBP-2 movements were generated using the Fiji plugin, KymographClear 2.0a (Mangeol et al., 2016). A 2-pixel width line was drawn to create the kymographs, and the number of lines, representing individual EBP-2 comets, and their direction were measured. Additionally, the velocities of each individual comet were calculated by measuring the length and height of each line.

Pioneer neuron dynamics: Nerve ring assembly was tracked by following the dynamics of the pioneer neurons labelled using the marker, P*lim-4*::mNG-PH and P*ceh-17*::mSca-I-PH. *lim-4* was expressed in 16 neurons whereas *ceh-17* expression was restricted to only SIAV pioneer neuron. The embryos expressing the markers were mounted on 2% agarose pad and movies of 12-13x 1μm z-stacks were acquired from the end of gastrulation, at 4 minute intervals for 4 hours using CFI60 Plan Apochromat lambda 60X (Nikon) objective (Figure 5). From the movies those embryos that displayed defects in the nerve ring were counted as defected. As defects in the nerve ring we counted embryos where the nerve ring did not form at all or nerve rings where some of the axons did not incorporate in the main axonal bundle. The standard error of the proportion was calculated according to: 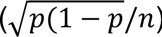, p equals the proportion of embryos with defects and n total number of embryos. To measure the positions of the pioneer neuron cell bodies in Figure 5E, the X,Y coordinates of the cell bodies of the neurons SIBV, SIAD, SMDD and SIAV were measured within the embryo in Fiji. These positions were then plotted, the Y axis represents the distance in the dorsoventral axis of the embryo, and the X axis represents the distance in the anterior-posterior axis of the embryo. Only one neuron of each neuronal pair was considered in the measurements.

### Quantification and statistical analysis

The data normality was assessed by using a Shapiro-Wilk test. Normally distributed data were compared with an unpaired t-test or a Mann-Whitney test (for comparison between two groups). The comparisons were done in GraphPad Prism and the stars ****, ***, ** and ns correspond to p<0.0001, p<0.001, p<0.01 and “not significant”, respectively.

## Supplemental information

**Figure S1:**
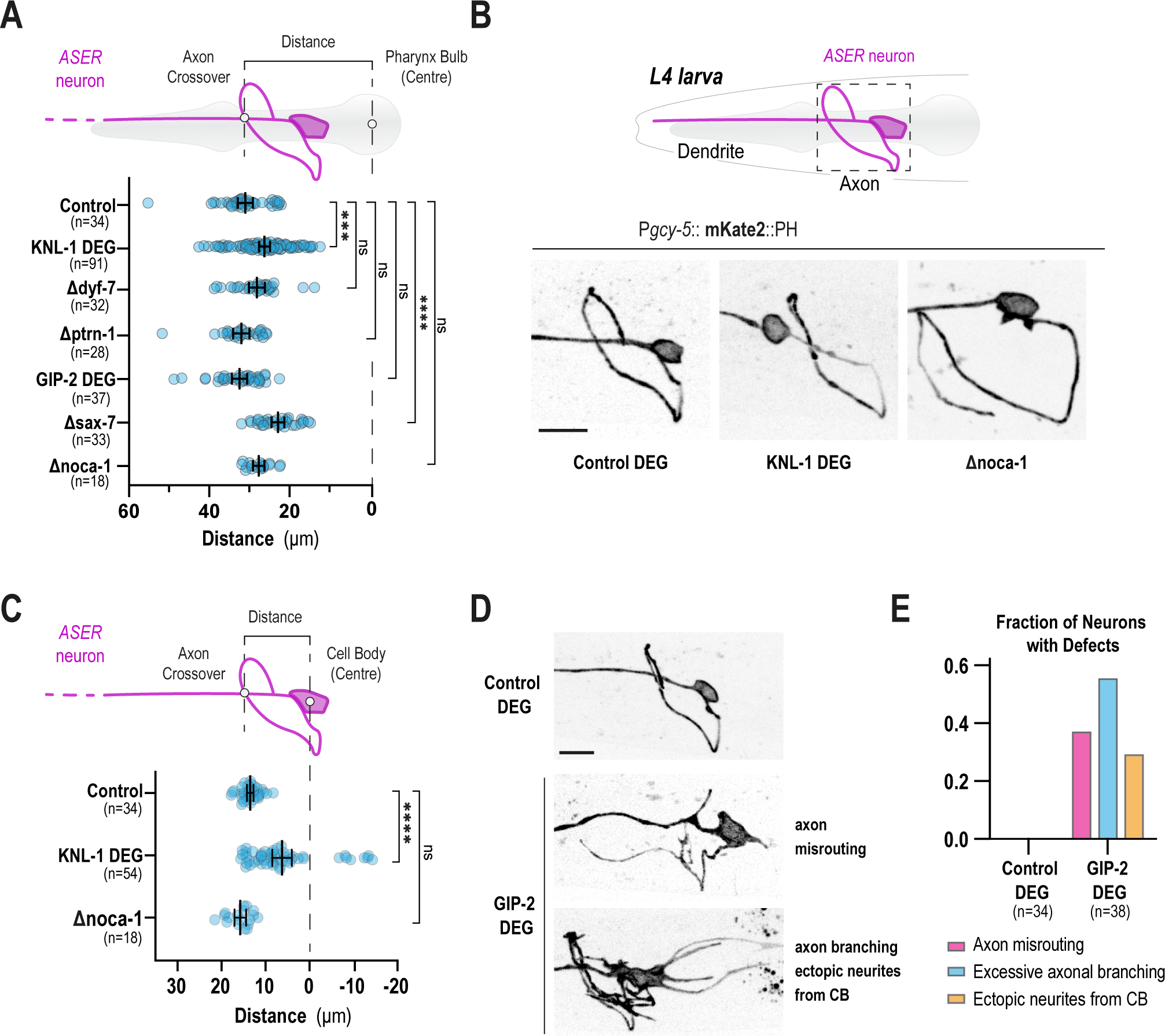
(A) Cartoon representation of the ASER neuron relatively to the terminal pharyngeal bulb of an L4 larva (top). Plot of distance between the pharynx bulb and the axon-dendrite intersect in the indicated conditions. n represents the number of animals. ***, **** and ns indicate p<0.001, p<0.0001 and non-significant, respectively. (B) Cartoon representation of the ASER neuron in an L4 larva (top). Crop-out images of the ASER axon and cell body in control, KNL-1 DEG and Δnoca-1. Scale bar, 10μm. (C) Illustrated schematic of the ASER neuron (top). Plot of distance between the centre of the ASER cell body and the axon-dendrite intersect in the indicated conditions (bottom). Control and KNL-1 DEG data are the same as in Figure 1D. n represents the number of animals. (D) Images of the range of ASER axonal phenotypes resulting from the post-mitotic degradation of GIP-2 using *dyf-7* driven GFP nanobody degrader system. GIP-2 degradation led to axon misrouting, where the axon did not follow the typical nerve ring trajectory, ectopic neurite formation, sprouting from the ASER cell body (CB), and ectopic axon branching. Scale bar, 10μm. (E) Quantification of the ASER axonal phenotypes of the control and GIP-2 DEG animals. n represents the number of neurons.

**Figure S2:**
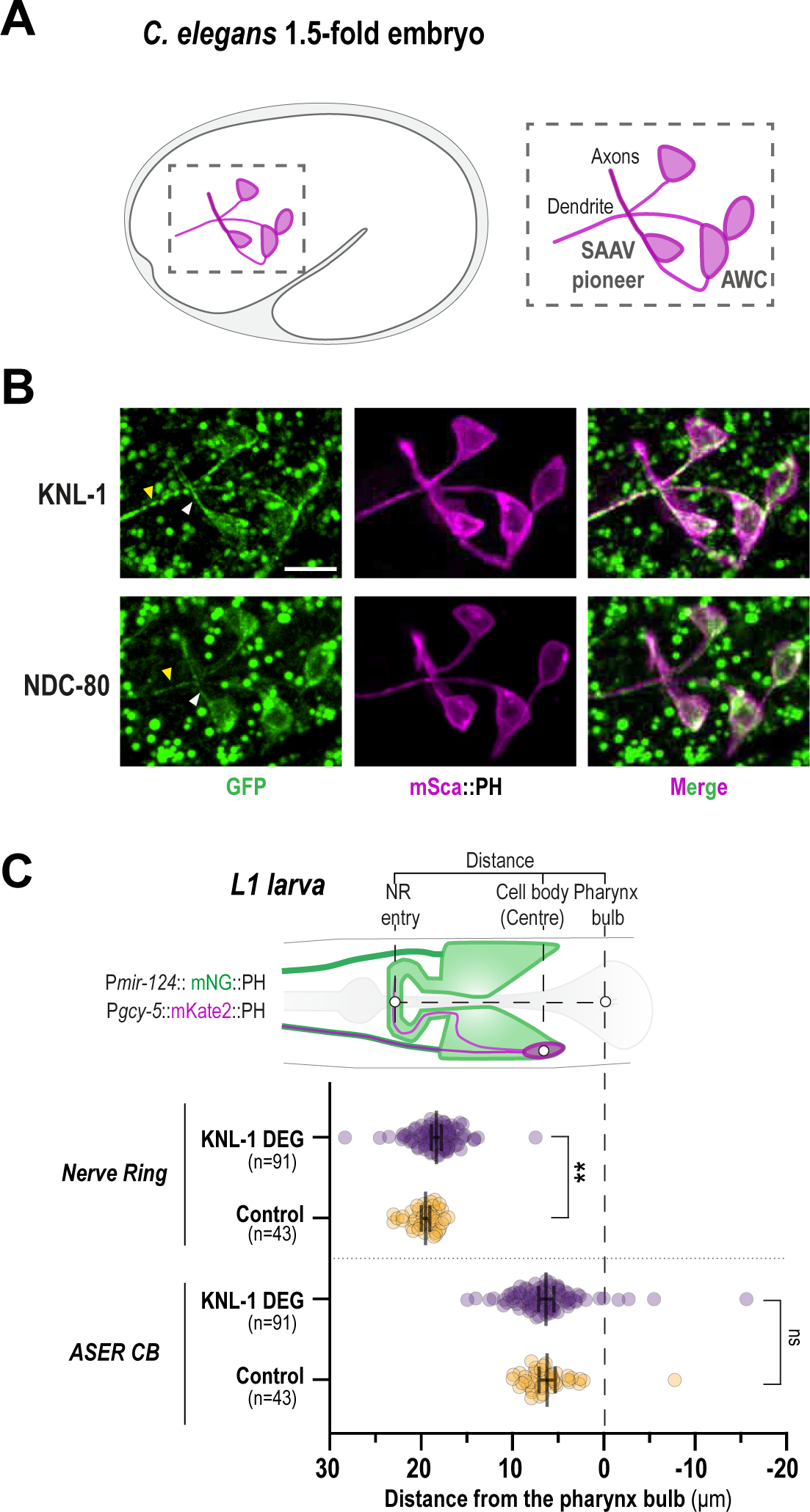
(A) Schematic illustration of a lateral view of a 1.5-fold C. elegans embryo highlighting P*hlh-16* expressing neurons. *hlh-16* is active in several neurons, including the AWC amphid neuron and the SAAV pioneer neuron, from bean stage of embryogenesis until L1 larva stage. The SAAV pioneer axon that forms the scaffold of the nerve ring has extended and integrated into the nerve ring structure. (B) Localization of GFP::KNL-1 and NDC-80::GFP in developing embryonic neurons, using a split-GFP system. Neuronal membranes are labelled with mSca-I::PH (magenta). White arrowheads indicate the presence of KNL-1 and NDC-80 in the axon of SAAV. Yellow arrowheads point out the presence of KNL-1 and NDC-80 in the AWC dendrite. Scale bar, 5μm. (C) Schematic illustrating the positions of ASER (magenta) and the P*mir-124* expressing sensory neurons (green) (top). Two different measurements were quantified from the images: distance of the pharynx bulb from the nerve ring (middle) and from the ASER cell body (bottom). The distances were measured as represented in the schematic. Compared to the control, the ASER cell body maintained the same position, while the nerve ring was misplaced anteriorly, relatively to the pharynx bulb, in KNL-1 DEG animals. n indicates the number of animals. Error bars denote 95% confidence interval. ** and ns indicate p<0.01 and non-significant, respectively.

**Figure S3:**
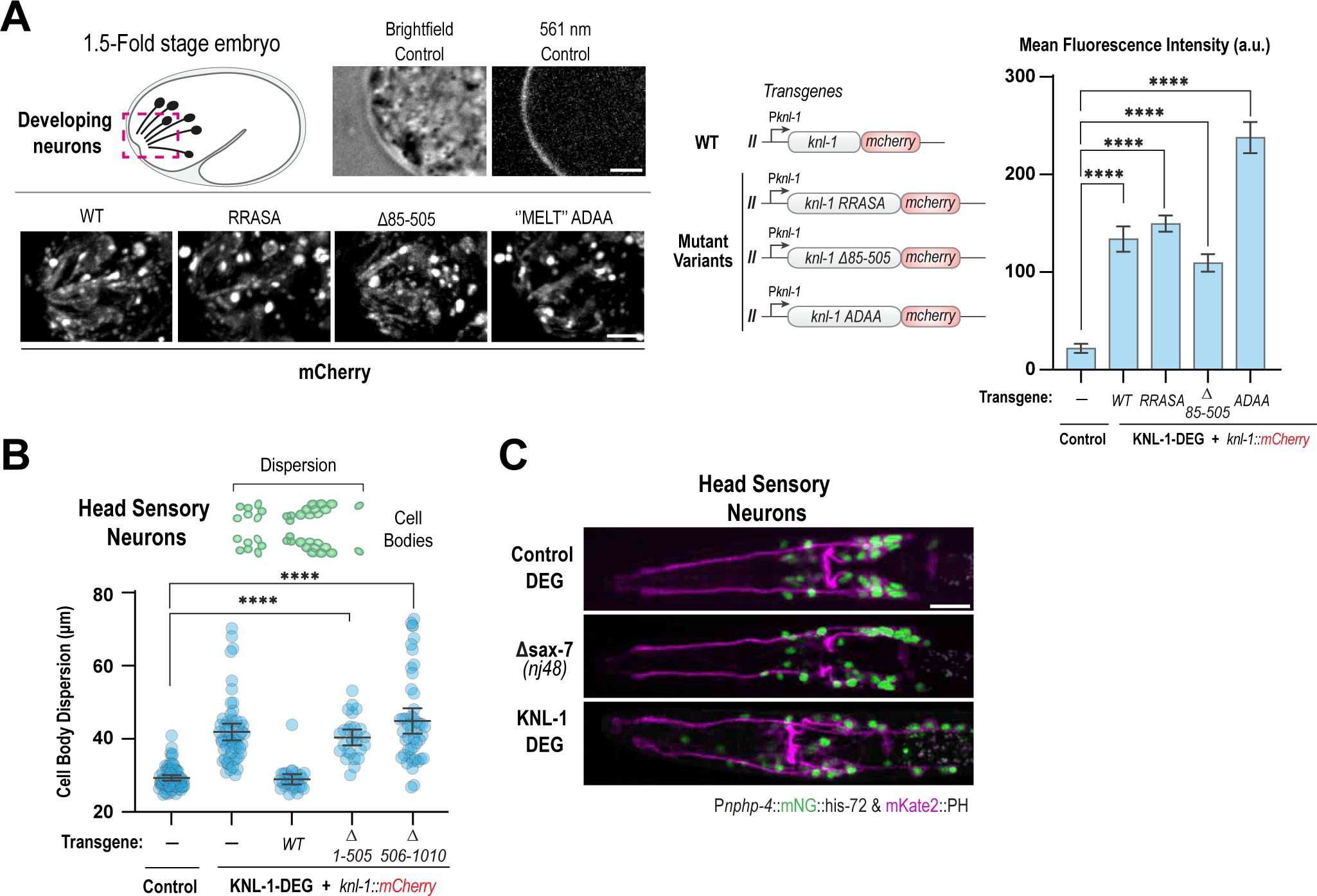
(A) Localization of KNL-1 variants in the developing dendrites of the sensory neurons in the 1.5-fold stage of embryogenesis, using the gene replacement system. Cartoon representing the developing dendrites in the embryo (top-left). The images at the top show a region of the embryo head of the control using Brightfield and fluorescence at 561nm. In the control there is no KNL-1::mCherry enriched in the dendrites. In the full length KNL-1 rescue (WT) KNL-1 is found in the dendrites. Similar localization pattern was observed in the expression of the KNL-1 mutant alleles (RRASA, Δ85-505, ‘’MELT’’ ADAA) in the background of the KNL-1 GFP degrader. Scale bar, 3μm. The schematic in the middle illustrates the various KNL-1 transgenes used in this assay. The different transgenes are expressed under *knl-1* promoter and they are fused to mCherry. The graph on the right represents the quantification of the mean fluorescence intensity of KNL-1::mCherry in the dendrites in the indicated conditions. Error bars denote 95% confidence interval. **** indicates p<0.0001. (B) Quantification of sensory neuron cell body dispersion in the indicated conditions. The gene replacement system was used to express two alleles of KNL-1: one lacking the N-terminal signalling hub (Δ1-505), the other lacking the C terminal Ndc-80 complex recruitment module (Δ506-1010). Deletion of either region significantly impacted the dispersion of head sensory neuron cell bodies. The effect of C-terminal deletion appears more pronounced based on the distribution of the values. n represents the number of animals. Control and KNL-1 WT and DEG data are the same as in Figure 3E. **** indicates p<0.0001. (C) Images of head sensory neuron nuclei (green) and plasma membranes (magenta) for the indicated conditions. Images of Control and KNL-1 DEG are same as Figure 3D. Scale bar,10 μ.

**Figure S4:**
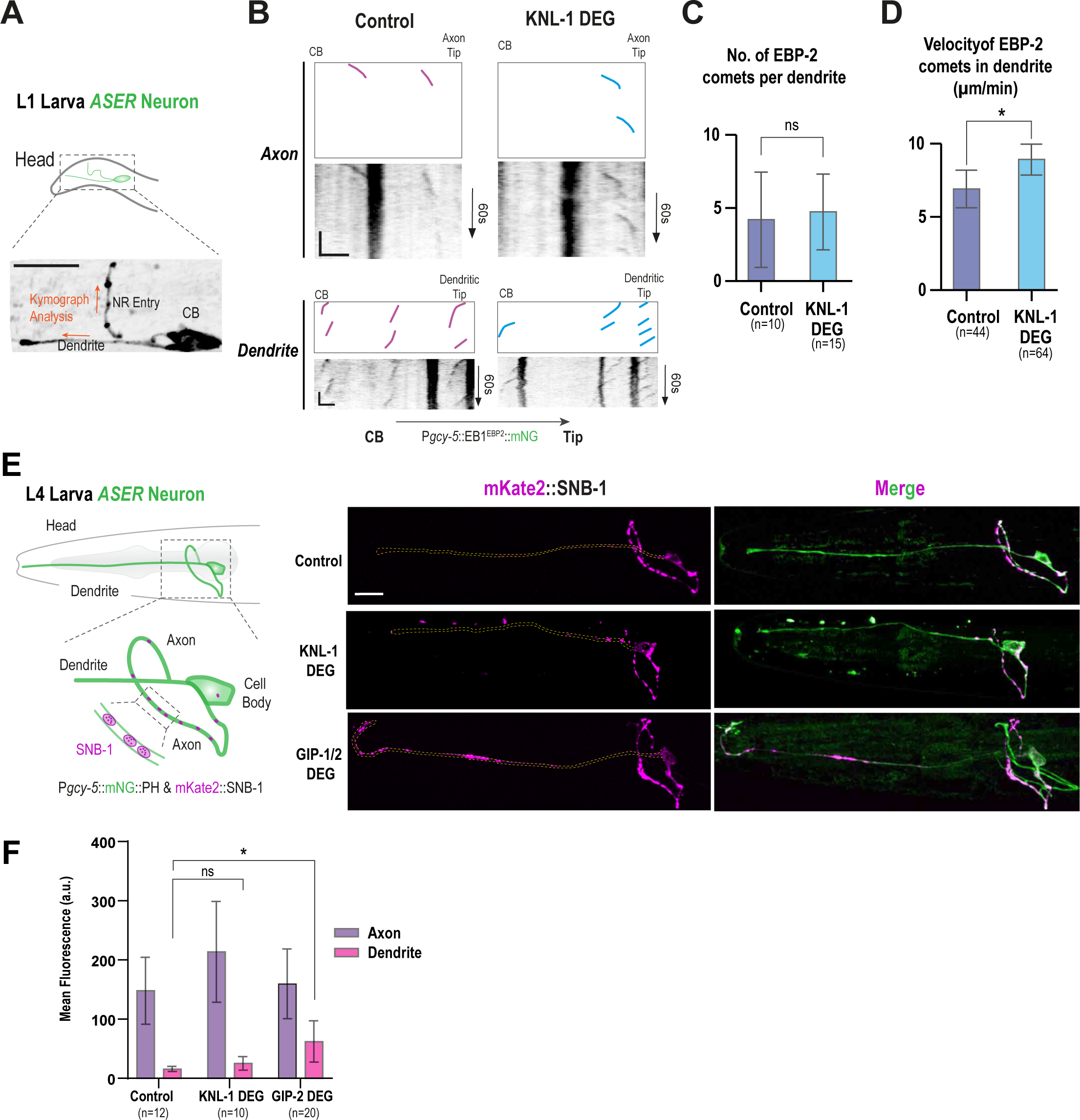
(A) EB1^EBP-2^::mNG dynamics in the ASER axon and dendrite. The image is a maximum intensity projection of a single time frame of EB1^EBP-2^::mNG movie. Scale bar, 10μm. (B) Kymographs of EB1^EBP-2^::mNG puncta in axons and dendrites of control and KNL-1 DEG animals. Axon kymographs were generated for the last section of the axon that enters the nerve ring, while for the dendrite, the kymographs were generated from the whole dendritic region. The direction of the kymograph is indicated by the orange arrow in (A). Scale bar, 2μm (horizontal), 15s (vertical). (C-D) Quantification of the number and velocity of comets per dendrite. Error bars denote 95% confidence interval. n represents the number of dendrites. * and ns indicate p<0.05 and non-significant, respectively. (E) Schematic showing the localization of synaptic marker SNB-1 in the ASER axon. To visualize ASER neuron morphology and SNB-1 protein localization, the transgenes P*gcy-5*::mNeonGreen and P*gcy 5*::mKate2::SNB-1 were utilized, respectively. The images on the right depict the localization of synaptic marker SNB-1 (magenta) in the axon and dendrite (highlighted with yellow dashed line) of the ASER (green) in control, KNL-1 DEG and GIP-2 DEG. SNB-1 puncta are found ectopically in the dendrite of the ASER post GIP-1/2 degradation, and not in the dendrites of control and KNL-1 DEG. Scale bar,10 μm. (E) Quantification of SNB-1 signal in the axon and dendrites, in the indicated conditions. Control and KNL-1 DEG data are the same as in Figure 4G. Error bars denote 95% confidence interval. n represents the number of animals. * and ns indicate p<0.05 and non-significant, respectively.

**Figure S5:**
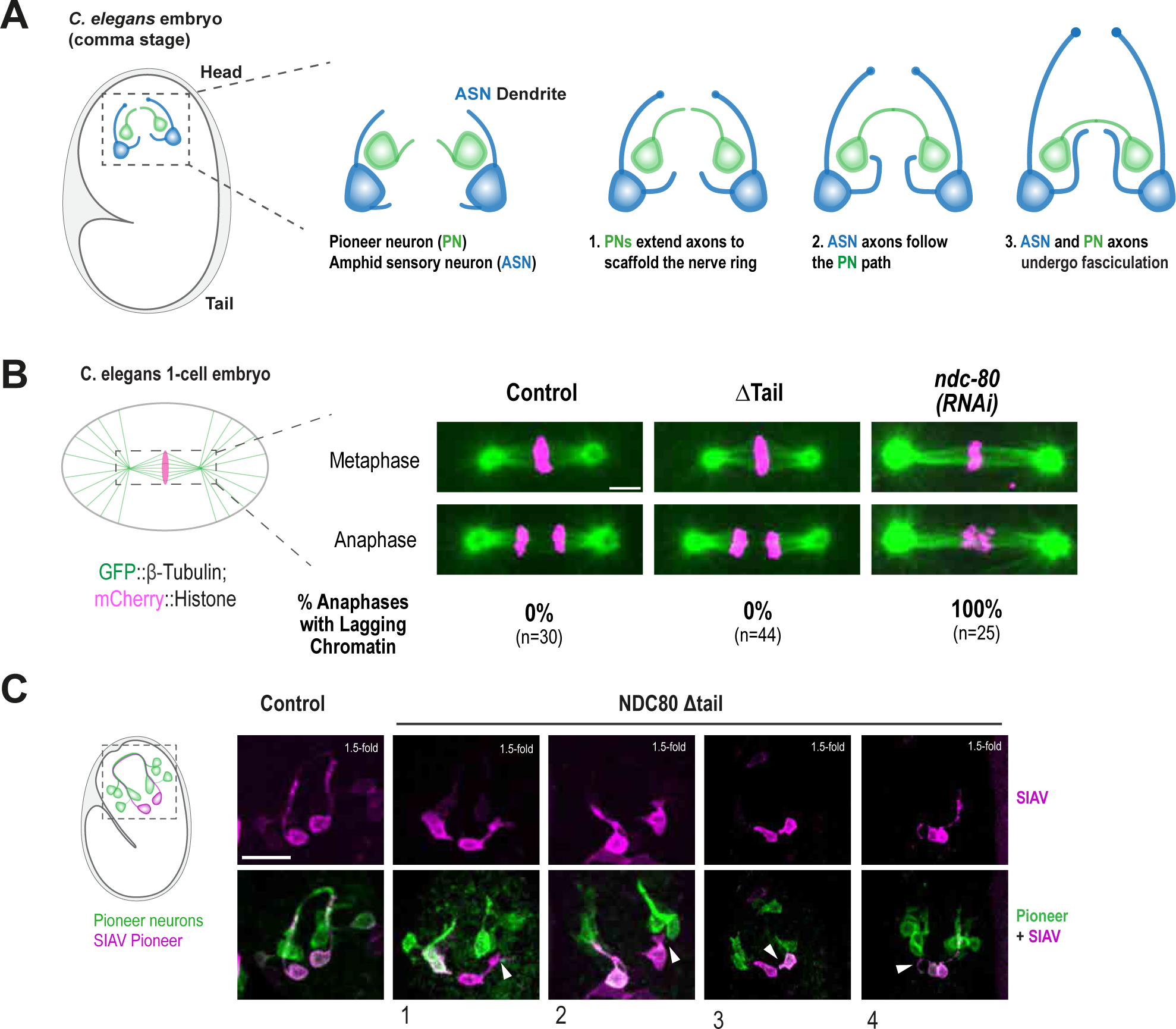
(A) Schematic of the nerve ring assembly in a C. elegans embryo. At comma stage, pioneer neuron axons extend to create the ring-shaped nerve ring scaffold. Subsequently, the axons of follower neurons, including the amphid sensory neurons, track the pioneer axons and undergo fasciculation to form the densely interconnected axonal structure in the nerve ring (Rapti et al. 2017; Moyle et al. 2021). (B) Schematic illustration of single-cell *C. elegans* embryo undergoing mitotic metaphase (left). Microtubules are labelled using GFP::TBB-2 (green) and the chromosomes are labelled using mCherry::H2B (magenta). Images on the right show the spindle and chromosomes during metaphase and anaphase of mitosis in the indicated conditions. The percentage of anaphases with lagging chromosomes is shown at the bottom of each panel. Upon NDC-80 depletion from the embryos through dsRNA mediated silencing, the chromosomes fail to align properly, resulting in lagging chromosomes during anaphase. In contrast, the NDC 80 ΔTail deletion did not show alignment defects or chromosome missegregation, suggesting that chromosome segregation was occurring normally in this condition. Scale bar, 5μm. (C) Schematic representation of the lateral view of a *C. elegans* embryo at 1.5-fold stage highlighting the pioneer neurons that form the nerve ring scaffold (left). Pioneer neurons were labelled using P*lim-4*::mNG::PH; P*ceh-17*::mSca::PH expression was restricted to SIAV neuron. Such labelling enabled tracking of the entry of SIAV axon into the pioneer axon bundle. The images display proper nerve ring assembly in the control 1.5-fold embryo alongside examples of SIAV (magenta) of axon bundling failures and inability to incorporate into the nerve ring scaffold (green). Examples 1,2 and 4 show cases where the SIAV axons are initially guided correctly towards the nerve ring but subsequently fail to extend (white arrowheads). In example 3 the SIAV axons fail to extend. Scale bar, 10μm.

## Supplementary methods

### RNA-mediated Interference

The RNAi against ndc-80 was performed by microinjection. Double-stranded RNAs were generated as described (Oegema et al., 2001) using DNA templates prepared by PCR-amplifying regions using the oligonucleotides:

5‘-GATGACAAGTACATTCAGAGATTATACAAATGATC-3’

5‘-GTGGTTCAAGATTCATTTGAATATTAAGTCCACTG-3’
and N2 genomic DNA as the template. L4 hermaphrodites were injected with dsRNA and incubated at 20°C for 36–46 h before dissection and imaging of their embryos.

### Fluorescence microscopy and analysis

To visualize the expression of GFP::KNL-1 and GFP::NDC-80 **(Figure S2)** in embryonic neurons, a split gfp system linked to mScarlet::PH under the *hlh-16* promoter was utilized. Neurons were initially identified based on mScarlet::PH marker expression, and 15×0.5μm z-stacks were acquired to capture all the neurons.

**Table S1:** *C. elegans* strains used in the study.

| GENOTYPE | SOURCE | IDENTIFIER | FIGURE |
| --- | --- | --- | --- |
| *C. elegans* N2 Bristol | Caenorhabditis Genetics Center | N2 |  |
| *lad-1(nj48)* | PMID: 15775964 | IK0441 | **1 & 3 & S1 & S3** |
| *ItSi220[pOD1249/pSW077; Pmex-5::GFP-tbb-2-operon-linker-mCherry-his-11; cb-unc-119(+)]I* | PMID: 26371552 | OD868 | **S5** |
| *ItSi1038 [pDC344; Pgcy-5::mKate-2::PH_unc-54_3'UTR; cb-unc-119(+)]II; unc-119(ed3)III* | PMID: 30827898 | OD3242 | **1** |
| *ItSi1016[pDC337; Pdyf-7::vhhGFP4::ZIF-1::dyf-7_3'UTR; cb-unc-119(+)]I #2; ItSi1038[pDC344; Pgcy-5::mKate-2::PH_unc-54_3'UTR; cb-unc-119(+)]II; unc-119(ed3)III; unc-119(ed3)III?; knl-1(It53[knl-1::GFP::tev::loxP::3xFlag])III* | PMID: 30827898 | OD3247 | **1 & S1** |
| *ItSi1016[pDC337; Pdyf-7::vhhGFP4::ZIF-1::dyf-7_3'UTR; cb-unc-119(+)]I; ItSi1038 [pDC344; Pgcy-5::mKate-2::PH_unc-54_3'UTR; cb-unc-119(+)]II ItSi1016[pDC337; Pdyf-7::vhhGFP4::ZIF-1::dyf-7_3'UTR; cb-unc-119(+)] knl-1(It53[knl-1::GFP::tev::loxP::3xFlag])III; unc-119(ed3)III ?* | PMID: 30827898 | OD3249 | **1 & S1** |
| *ItSi1038 [pDC344; Pgcy-5::mKate-2::PH_unc-54_3'UTR; cb-unc-119(+)]II ; unc-119(ed3)III?; dyf-7 (it60)X* | PMID: 30827898 | OD3291 | **1 & S1** |
| *ItSi1016[pDC337; Pdyf-7::vhhGFP4::ZIF-1::dyf-7_3'UTR; cb-unc-119(+)]#2 I] ; ItSi1[pOD809/pJE110; Pknl-1::KNL-1reencoded::RFP; cb-unc-119(+)]II; knl-1(It53[knl-1::GFP::tev::loxP::3xFlag]) III* | PMID: 30827898 | OD3346 | **3 & S3** |
| *ItSi1016[pDC337; Pdyf-7::vhhGFP4::ZIF-1::dyf-7_3'UTR; cb-unc-119(+)]#2 I] ; ItSi13[pOD858/pJE123; Pknl-1::KNL-1aa506-1010::mCherry; cb-unc-119(+)]II; knl-1(It53[knl-1::GFP::tev::loxP::3xFlag]) III* | PMID: 30827898 | OD3404 | **S3** |
| *unc-119(ed3)III; ItSi1174[oxTi365; pDC591; Pnphp-4::mNeonGreen-his-72:tbb-2_3'UTR;;gpd-2/3 operon linker-mKate2-PH:unc-34_3'UTR]V* | PMID: 30827898 | OD3919 |  |
| *ItSi1016[pDC337; Pdyf-7::vhhGFP4::ZIF-1::dyf-7_3'UTR; cb-unc-119(+)]I; unc-119(ed3)III?; ItSi1174[oxTi365; pDC591; Pnphp-4::mNeonGreen-his-72:tbb-2_3'UTR;;gpd-2/3 operon linker-mKate2-PH:unc-34_3'UTR]V* | PMID: 30827898 | OD3924 | **3 & S3** |
| *ItSi1016[pDC337; Pdyf-7::vhhGFP4::ZIF-1::dyf-7_3'UTR; cb-unc-119(+)]I; knl-1(It53[knl-1::GFP::tev::loxP::3xFlag])III; ItSi1174[oxTi365; pDC591; Pnphp-4::mNeonGreen-his-72:tbb-2_3'UTR;;gpd-2/3 operon linker-mKate2-PH:unc-34_3'UTR]V* | PMID: 30827898 | OD3938 | **3 & S3** |
| ItSi1016[pDC337; Pdyf-7::vhhGFP4::ZIF-1::dyf-7_3'UTR; cb-unc-119(+)]I ;<br>ItSi120[[pDC170;Pndc-80:NDC-80 reencoded; cb-unc-119(+)]II #3; unc-119(ed3)III? ; ndc-80(It54[ndc-80::GFP::tev::loxP::3xFlag])IV; Pnphp-4::mNeonGreen-his-72:tbb-2_3'UTR;;gpd-2/3 operon linker-mKate2-PH:unc-34_3'UTR]V | PMID: 30827898 | OD3952 | **3** |
| ItSi1016[pDC337; Pdyf-7::vhhGFP4::ZIF-1::dyf-7_3'UTR; cb-unc-119(+)]I ;<br>ItSi121[pDC175;Pndc-80:NDC-80 (Mutant D1-59) reencoded; cb-unc-119(+)]II #1; unc-119(ed3)III? ; ndc-80(It54[ndc-80::GFP::tev::loxP::3xFlag])IV; Pnphp-4::mNeonGreen-his-72:tbb-2_3'UTR;;gpd-2/3 operon linker-mKate2-PH:unc-34_3'UTR]V | PMID: 30827898 | OD3953 | **3** |
| ItSi1016[pDC337; Pdyf-7::vhhGFP4::ZIF-1::dyf-7_3'UTR; cb-unc-119(+)]I; ItSi711[pDC267;Pndc-80:NDC-80(66,96,100,125,144,155AAAAAA) reencoded; cb-unc-119(+)]II#1; unc-119(ed3)III? ; ndc-80(It54[ndc-80::GFP::tev::loxP::3xFlag])IV; ItSi1055[oxTi365;pDC344; Pnphp-4::mNeonGreen-his-72:tbb-2_3'UTR;;gpd-2/3 operon linker-mKate2-PH:unc-34_3'UTR]V | PMID: 30827898 | OD3954 | **3** |
| ItSi1016[pDC337; Pdyf-7::vhhGFP4::ZIF-1::dyf-7_3'UTR; cb-unc-119(+)] I ;<br>ItSi1[pOD809/pJE110; Pknl-1::KNL-1reencoded::RFP; cb-unc-119(+)]II; knl-1(It53[knl-1::GFP::tev::loxP::3xFlag]) III; ItSi1174[oxTi365; pDC591; Pnphp-4::mNeonGreen-his-72:tbb-2_3'UTR;;gpd-2/3 operon linker-mKate2-PH:unc-34_3'UTR]V | PMID: 30827898 | OD3977 | **3 & S3** |
| ItSi1016[pDC337; Pdyf-7::vhhGFP4::ZIF-1::dyf-7_3'UTR; cb-unc-119(+)]I ; unc-119(ed3)III? ; ndc-80(It54[ndc-80::GFP::tev::loxP::3xFlag])IV; ItSi1174[oxTi365; pDC591; Pnphp-4::mNeonGreen-his-72:tbb-2_3'UTR;;gpd-2/3 operon linker-mKate2-PH:unc-34_3'UTR]V | PMID: 30827898 | OD3998 | **3** |
| ItSi1016[pDC337; Pdyf-7::vhhGFP4::ZIF-1::dyf-7_3'UTR; cb-unc-119(+)]I; ItSi1191[pDC603; Pgcy-5::mNeonGreen-PH::tbb-3'UTR; Pgcy-5::mKate2-snb-1::snb-1_3'UTR; cb-unc-119(+)] II; unc-119(ed3) III? | PMID: 30827898 | OD4011 | **4 & S4** |
| ItSi1016[pDC337; Pdyf-7::vhhGFP4::ZIF-1::dyf-7_3'UTR; cb-unc-119(+)] I ; ItSi1191[pDC603; Pgcy-5::mNeonGreen-PH::tbb-3'UTR; Pgcy-5::mKate2-snb-1::snb-1_3'UTR; cb-unc-119(+)] II; unc-119(ed3) III; knl-1(It53[knl-1::GFP::tev::loxP::3xFlag])III | PMID: 30827898 | OD4012 | **4 & S4** |
| ItSi1016[pDC337; Pdyf-7::vhhGFP4::ZIF-1::dyf-7_3'UTR; cb-unc-119(+)]#2 I ;ItSi13[pOD858/pJE123; Pknl-1::KNL-1aa506- | This study | DKC13 | **S3** |
| 1010::mCherry; cb-unc-119(+)]II; knl-1(lt53[knl-1::GFP::tev::loxP::3xFlag]) III; Pnphp-4::mNeonGreen-his-72:tbb-2_3'UTR;;gpd-2/3 operon linker-mKate2-PH:unc-54_3'UTR]V |  |  |  |
| ltSi1016[pDC337; Pdyf-7::vhhGFP4::ZIF-1::dyf-7_3'UTR; cb-unc-119(+)]#2 I; ltSi13[pOD858/pJE123; Pknl-1::KNL-1aa1-505::mCherry; cb-unc-119(+)]II; knl-1(lt53[knl-1::GFP::tev::loxP::3xFlag])III; Pnphp-4::mNeonGreen-his-72:tbb-2_3'UTR;;gpd-2/3 operon linker-mKate2-PH:unc-54_3'UTR]V | This study | DKC16 | **S3** |
| ltSi1016[pDC337; Pdyf-7::vhhGFP4::ZIF-1::dyf-7_3'UTR; cb-unc-119(+)]#2 I; ltSi9[pOD831/pJE120; Pknl-1::KNL-1reencoded(RRASAmutant)::RFP; cb-unc-119(+)]II; knl-1(lt53[knl-1::GFP::tev::loxP::3xFlag]) III | This study | DKC67 | **3 & S3** |
| ltSi1016[pDC337; Pdyf-7::vhhGFP4::ZIF-1::dyf-7_3'UTR; cb-unc-119(+)]#2 I; ltSi44[pOD1039/pJE170; Pknl-1::KNL-1reencoded(Mutant D85-505)::RFP; cb-unc-119(+)]II;knl-1(lt53[knl-1::GFP::tev::loxP::3xFlag])III | This study | DKC68 | **3 & S3** |
| ltSi1016[pDC337; Pdyf-7::vhhGFP4::ZIF-1::dyf-7_3'UTR; cb-unc-119(+)]#2 I; ltSi44[pOD1039/pJE170; Pknl-1::KNL-1reencoded(Mutant D85-505)::RFP; cb-unc-119(+)]II;knl-1(lt53[knl-1::GFP::tev::loxP::3xFlag]) III;Pnphp-4::mNeonGreen-his-72:tbb-2_3'UTR;;gpd-2/3 operon linker-mKate2-PH:unc-54_3'UTR]V | This study | DKC77 | **3 & S3** |
| ltSi1016[pDC337; Pdyf-7::vhhGFP4::ZIF-1::dyf-7_3'UTR; cb-unc-119(+)]#2 I; unc-119(ed3)III ?]; ltSi9[pOD831/pJE120; Pknl-1::KNL-1reencoded(RRASAmutant)::RFP; cb-unc-119(+)]II; knl-1(lt53[knl-1::GFP::tev::loxP::3xFlag]) III; Pnphp-4::mNeonGreen-his-72:tbb-2_3'UTR;;gpd-2/3 operon linker-mKate2-PH:unc-54_3'UTR]V | This study | DKC84 | **3 & S3** |
| ltSi1016[pDC337; Pdyf-7::vhhGFP4::ZIF-1::dyf-7_3'UTR; cb-unc-119(+)]#2 I; ltSi48[pOD1038/pJE169; Pknl-1::KNL-1reencoded(MELT repeats mutant)::RFP; cb-unc-119(+)]II; knl-1(lt53[knl-1::GFP::tev::loxP::3xFlag]) III | This study | DKC99 | **3 & S3** |
| ltSi1016[pDC337; Pdyf-7::vhhGFP4::ZIF-1::dyf-7_3'UTR; cb-unc-119(+)]#2 I; [Pknl-1::KNL-1 MELT deletion::RFP; cb-unc-119(+)]II; knl-1(lt53[knl-1::GFP::tev::loxP::3xFlag]) III; Pnphp-4::mNeonGreen-his-72:tbb-2_3'UTR;;gpd-2/3 operon linker-mKate2-PH:unc-54_3'UTR]V | This study | DKC105 | **3 & S3** |
| *ItSi1038 [pDC344; Pgcy-5::mKate-2::PH_unc-54_3'UTR; cb-unc-119(+)]II; unc-119(ed3)III ?; ptrn-1(It1::cb-unc-119+)X* | This study | DKC112 | **1 & S1** |
| *ItSi1016[pDC337; Pdyf-7::vhhGFP4::ZIF-1::dyf-7_3'UTR; cb-unc-119(+)]I; gip-2(It19[gip-2::GFP]::loxP::cb-unc-119(+))::loxP)I; ItSi1019[pDC353; Pdyf-7::vhhGFP4::ZIF-1::dyf-7_3'UTR; cb-unc-119(+)]II ; unc-119(ed3)III ?; ItSi1055[oxTi365;[pDC344; Pgcy-5::mKate-2::PH_unc-54_3'UTR; cb-unc-119(+)]V* | This study | DKC129 | **S1** |
| *ndc-80(tm5271)/ IV; ItSi1016[pDC337; Pdyf-7::vhhGFP4::ZIF-1::dyf-7_3'UTR; cb-unc-119(+)]I; ItSi122[pDC178;Pndc-80:NDC-80 (8,18,44,51AAAA) reencoded; cb-unc-119(+)]II #1; unc-119(ed3)III?; ItSi1174[oxTi365; pDC591; Pnphp-4::mNeonGreen-his-72:tbb-2_3'UTR;;gpd-2/3 operon linker-mKate2-PH:unc-54_3'UTR]V* | This study | DKC160 | **3** |
| *ItSi1038 [pDC344; Pgcy-5::mKate-2::PH_unc-54_3'UTR; cb-unc-119(+)]II; unc-119(ed3)III?]; noca-1(ok3692)V/nT1[qIs51](IV;V)* | This study | DKC162 | **S1** |
| *ndc-80(dha60; NDC-80 deltaTail)IV* | This study | DKC400 | **5 & S5** |
| *ItSi220[pOD1249/pSW077; Pmex-5::GFP-tbb-2-operon-linker-mCherry-his-11; cb-unc-119(+)]I; ndc-80(dha60; NDC-80 deltaTail)IV* | This study | DKC402 | **S5** |
| *ItSi1172[pDC591; Pnphp-4::mNeonGreen-his-72:tbb-2_3'UTR;;gpd-2/3 operon linker-mKate2-PH:unc-54_3'UTR]II; unc-119(ed3)III; sax-7(nj48) IV* | This study | DKC454 | **3 & S3** |
| *unc-119(ed3)III; sax-7(nj48) IV; ItSi1055[oxTi365;[pDC344; Pgcy-5::mKate-2::PH_unc-54_3'UTR; cb-unc-119(+)]V* | This study | DKC559 | **1 & S1** |
| *knl-1(dha121(AID::darkg>f>p:::KNL-1) )III* | This study | DKC614 | **4 & S4** |
| *knl-1(dha130; HA::7XGFP11::knl-1)III* | This study | DKC693 | **2 & 5 & S2** |
| *dhaSi115[pDC1034; h1h-16::splitGFP1-10::rab-3_3'UTR ;;gpd-2 3operon::mSca-I-PH::unc-54_3'UTR; cb-unc-119(+)]II; knl-1(dha130; HA::7XGFP11::knl-1)III* | This study | DKC896 | **2 & 5 & S2** |
| *dhaSi114[pDC717; lim-4 pro::mNG::PH::unc-54_3'UTR]II; dhaSi73[pDC725; ceh-17 pro::mSca-PH::unc-54_3'UTR; cb-unc-119(+)] IV; unc-119(ed3)III* | This study | DKC1156 | **5 & S5** |
| *ItSi1016[pDC337; Pdyf-7::vhhGFP4::ZIF-1::dyf-7_3'UTR; cb-unc-119(+)]I #2 ; unc-119(ed3)III?; dhaSi148[pDC897;Pmir-124::mNG-PH::unc-54_3'UTR]II; knl-1(dha121(AID::darkg>f>p:::KNL-1) )III; unc-119(ed3)III; ItSi1055[oxTi365;[pDC344; Pgcy-5::mKate-2::PH_unc-54_3'UTR; cb-unc-119(+)]V* | This study | DKC1185 | **2 & S2** |
| *ItSi1016[pDC337; Pdyf-7::vhhGFP4::ZIF-1::dyf-7_3'UTR; cb-unc-119(+)]I #2 ; dhaSi148[pDC897;Pmir-124::mNG-PH::unc-54_3'UTR]II; unc-119(ed3)III; ItSi1055[oxTi365;pDC344; Pgcy-5::mKate-2::PH_ unc-54_3'UTR; cb-unc-119(+)]V* | This study | DKC1188 | **2 & S2** |
| *dhaSi114[pDC717; lim-4 pro::mNG::PH::unc-54_3'UTR]II; unc-119(ed3)III; dhaSi73[pDC725; ceh-17 pro::mSca-PH::unc-54_3'UTR; cb-unc-119(+)] IV; dhaSi114[pDC717; lim-4 pro::mNG::PH::unc-54_3'UTR]II; ndc-80(dha60; NDC-80 deltaTail)IV* | This study | DKC1194 | **5 & S5** |
| *ItSi1016[pDC337; Pdyf-7::vhhGFP4::ZIF-1::dyf-7_3'UTR; cb-unc-119(+)]I #2 ; dhaSi305[pDC716; Pgcy-5::mNG::ebp-2::tbb-2_3'UTR::rlaop::mScaPH::unc-54_3'UTR; cb-unc-119(+)]II; unc-119(ed3)III?; knl-1(dha121(AID::darkg>f>p:::KNL-1) )III* | This study | DKC1211 | **4 & S4** |
| *ItSi1016[pDC337; Pdyf-7::vhhGFP4::ZIF-1::dyf-7_3'UTR; cb-unc-119(+)]I #2 dhaSi305[pDC716; Pgcy-5::mNG::ebp-2::tbb-2_3'UTR::rlaop::mScaPH::unc-54_3'UTR; cb-unc-119(+)]II;; unc-119(ed3)III?* | This study | DKC1213 | **4 & S4** |
| *dhaSi115[pDC1034; h1h-16::splitGFP1-10::rab-3_3'UTR ::gpd-2/3operon::mSca-I-PH::unc-54_3'UTR; cb-unc-119(+)]II; ndc-80[It126 (ndc-80::7XGFP-11)] IV* | This study | DKC1234 | **5 & S2** |
| *ItSi1016[pDC337; Pdyf-7::vhhGFP4::ZIF-1::dyf-7_3'UTR; cb-unc-119(+)]I; gip-2(It19[gip-2::GFP]::loxP::cb-unc-119(+)::loxP)I; ItSi1191[pDC603; Pgcy-5::mNeonGreen-PH::tbb-3'UTR; Pgcy-5::mKate2-snb-1::snb-1_3'UTR; cb-unc-119(+)] II; unc-119(ed3)III?; gip-1(wow3[gfp::gip-1])III* | This study | DKC1321 | **S4** |

**Table S2:**
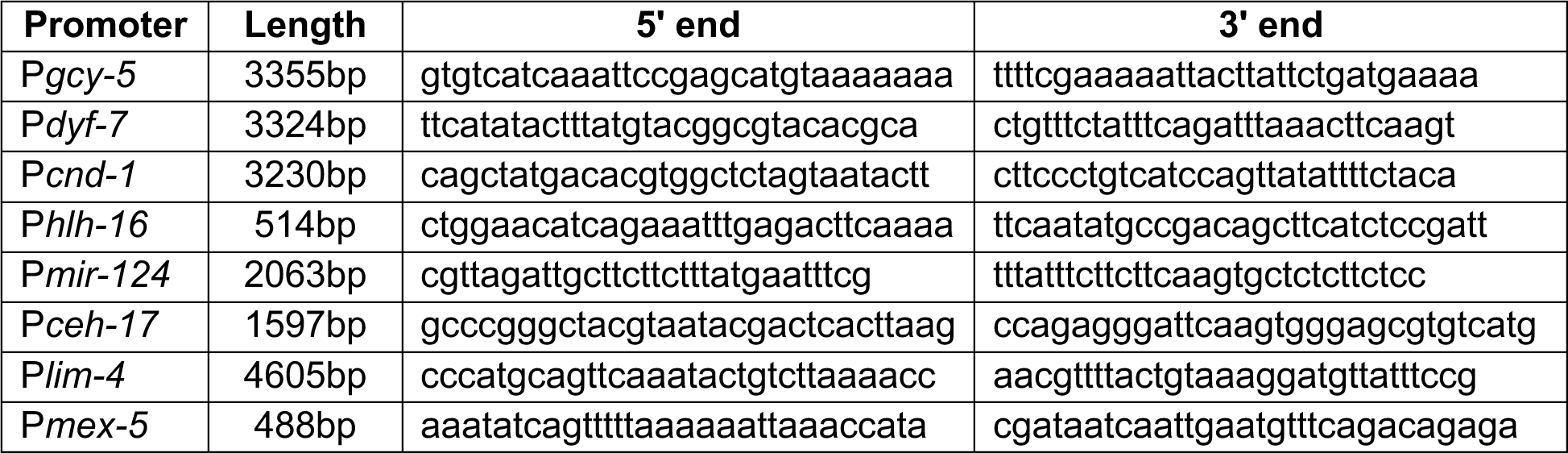
Sequences of regulatory elements.

**Table S3:**
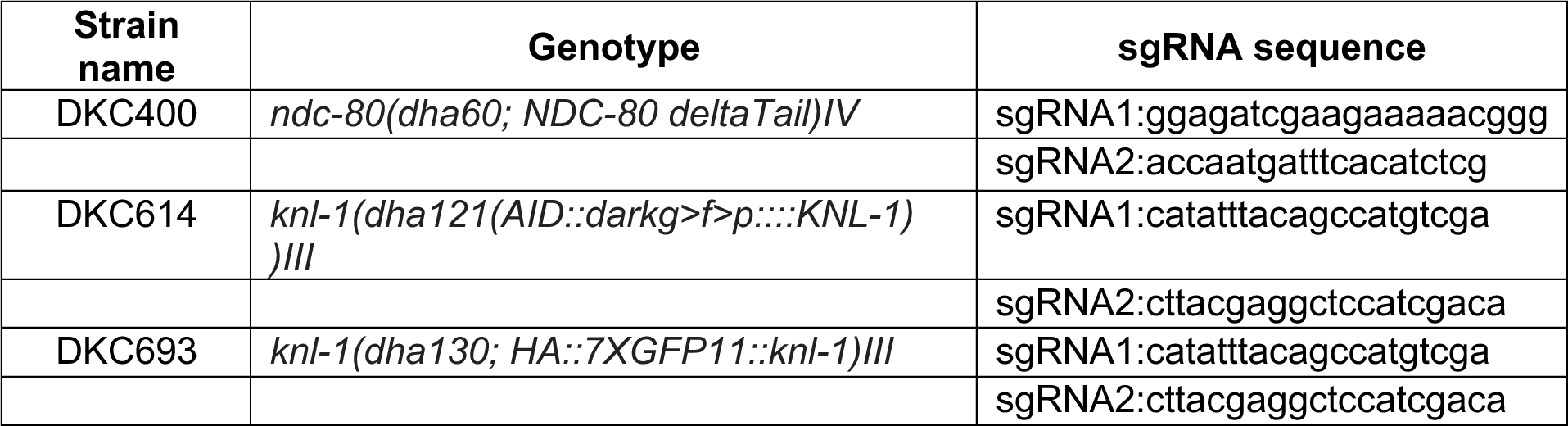
CRISPR-Cas9 loci & sgRNA sequences used for strain generation.

